# One cannot have it all: trading-off ecosystem services and biodiversity bundles in landscape connectivity restoration

**DOI:** 10.1101/2024.11.27.624888

**Authors:** Neyret Margot, Richards Daniel, Prima Marie-Caroline, Thomas R. Etherington, Lavorel Sandra

## Abstract

Countering the impacts of habitat loss and fragmentation on ecosystems requires complementing conservation areas with Other Effective area-based Conservation Measures within landscapes to promote biodiversity and multiple ecosystem services (ES). However, critical knowledge gaps persist in where and how natural elements should be restored to improve landscape connectivity to simultaneously support, and reduce trade-offs between biodiversity and ES. In virtual landscape experiments that allow exploring the effects of spatial pattern systematically, we generated alternative landscape restoration scenarios aimed at fostering ecological connectivity. Scenarios varied in the location and size of restored areas complementing existing natural areas. We analysed the impact of these scenarios on four bundles representing distinct priorities of target ES and biodiversity-related values. As expected, all bundles were favoured by increasing restored area in the landscape, but they were promoted by different spatial configurations. Restoration scenarios that fostered high aggregation of natural habitats promoted biodiversity and cultural value-related bundles, while smaller natural elements dispersed throughout the landscape were more beneficial for the sustainable production and climate adaptation bundles. These contrasts were most pronounced at low restoration efforts, where landscape configuration had greatest impacts on biodiversity and ecosystem processes. Effective spatial planning of restoration initiatives within landscapes should consider these trade-offs, along with context-specific constraints, when prioritizing areas for restoration or conservation. Our findings contribute to a more comprehensive understanding of how protected and restored areas can be integrated within landscapes to jointly support connectivity for both biodiversity and people.

**Highlights:**

- Virtual landscape restoration options effects on four ecosystem services and biodiversity bundles
- High aggregation of restored elements promoted *biodiversity* and *cultural value*
- Low aggregation promoted *sustainable production* and *climate adaptation*
- These contrasts were most important at low restoration targets
- These results highlight the importance of configuration trade-offs in restoration planning

## Introduction

The adverse effects of human-induced land conversion on biodiversity and the ecosystem services (ES) it underpins has been extensively documented (IPBES, 2019). In particular, the landscape- and regional-scale fragmentation of natural habitats is one of the key drivers of biodiversity loss (Haddad *et al*., 2015). Due to increasing recognition that human good quality of life depend on biodiverse and healthy ecosystems, there are growing calls to protect and restore ecosystem integrity including connectivity as one of its critical dimensions (Nicholson *et al*., 2021; CBD, 2022). Among the approaches proposed by the Global Biodiversity Framework, Other Effective area-based Conservation Measures offer a complementary strategy to protected areas. In support of their implementation, understanding how integrated conservation, including high-quality habitat under protection and restoration within production landscapes (i.e., landscapes that support both biodiversity and the provision of goods and services for humans (IPSI Secretariat, 2018)), can improve ecological connectivity for multiple taxa and a diversity of ES is a major challenge.

Much research has aimed to assess, maintain and restore landscape connectivity for biodiversity (Arroyo-Rodríguez *et al*., 2020) and ES (Grass *et al*., 2019). Complementary to structural connectivity, defined in terms of the spatial extent and distribution of habitat patches, which reflects landscape composition and configuration, landscape functional connectivity describes the ability of different biota to move in a landscape depending on their characteristics such as dispersal abilities (Tischendorf and Fahrig, 2000). Landscape connectivity thus varies across organisms, and its effects on ecosystems can be either positive, such as when it allows organisms to track their climatic niche (McGuire *et al*., 2016) or negative, when connectivity facilitates the movement of pests, diseases or invasive species (Fahrig, 2017). The effects of habitat aggregation (opposite of fragmentation) on connectivity are expected to be more pronounced when the availability of suitable habitats is low, as physical percolation between adjacent patches can be attained in non-random landscapes when habitat cover reaches 30-40% (Gardner *et al*., 1992).

Landscape connectivity also matters for ES. ES rely on the presence and activity of ES-providing biota (Luck et al., 2009) and on specific biophysical processes like flows of nutrients and water (affecting ES such as water quality, Brauman et al., 2007) or soil stabilisation (relevant for e.g. landslide prevention, Baró et al., 2017)), many of which are sensitive to landscape configuration (Qiu, 2019). Landscape configuration can have contrasted effects depending on the processes underlying each ES. For instance, habitat aggregation positively impacts perceived landscape visual quality (Wartmann *et al*., 2021). Pollination and pest control, on the other hand, tend to benefit from higher flow of ES providers to agricultural areas in complex patchy landscapes (Mitchell *et al*., 2015; Sirami *et al*., 2019). These varying responses make it difficult to predict the effects of the configuration of restored habitats in the landscape on multiple ES.

Despite their interconnections, it is common for research and policy to treat connectivity restoration for biodiversity and ES as separate issues, which challenges the development of integrated planning for restoration of landscape connectivity. Exceptions to this include studies investigating integrated landscape management options to promote biodiversity and single services such as agricultural production (Rey Benayas and Bullock, 2012; Grass *et al*., 2019), invasive species control (Buchholtz et al. 2023), pollination (López-Cubillos *et al*., 2023) and pest control (Kärvemo *et al*., 2017). However, these studies usually focus on few ES and taxonomic or functional groups. Others have developed approaches to prioritise restoration areas that optimise ecosystem services and/or connectivity, using tools such as circuit theory to identify “pinch points” for animal movement in target landscapes (McRae *et al*., 2008). However, the outcomes of such approaches are case-specific and an integrated understanding of how different scenarios of restoration impact connectivity for multiple biota and the resulting consequences for ES is lacking. In particular, which landscape configurations promote which land use objectives, such as culturally-motivated biodiversity conservation or sustainable agricultural production, is unclear.

The effects of landscape connectivity and fragmentation on biodiversity and ES have been studied through observational studies (Valdés *et al*., 2020) or experimental manipulation of landscape configuration (Damschen *et al*., 2019). These approaches are crucial for assessing local ES responses and their underlying mechanisms. Yet, they are constrained by local conditions and restrict the scope of investigation to existing landscapes. In response to these limitations, several studies have turned to virtual landscapes as a means of examining systematic variations in landscape composition and configuration (Mitchell, Bennett and Gonzalez, 2015; Lavorel *et al*., 2022), paving the way towards a more generalisable understanding of the impacts of landscape structure on multiple ES.

In this study, we investigate the impact of landscape restoration scenarios on multiple ES and dispersal kernel-based landscape connectivity for generic species groups in virtual landscape experiments. We selected archetypal connectivity groups (hereafter: connectivity groups) to represent contrasted sensitivities to land cover and fragmentation, in terms of habitat suitability, dispersal abilities, and minimum habitat size. The restoration scenarios varied in the extent and type of the natural elements restored (5%, 10%, or 20% including hedges, riparian strips, and forest patches) and their spatial configuration. Three configuration treatments were chosen to generate contrasted patterns of aggregation of natural elements: restoring parcels at random, along least-cost path corridors between existing natural areas (Etherington, 2016), or in large patches most distant from existing natural areas. We compared these in a factorial design to assess their impact on nine ES and six connectivity groups. Scenario outcomes were analysed considering bundles representing different landscape objectives: sustainable agricultural production, climate change adaptation and mitigation, landscape cultural values, and biodiversity conservation. These bundles broadly represent ES demand bundles (Zoderer *et al*., 2019), but also included connectivity groups in addition to ecosystem services, as different groups were considered relevant for each objective. We hypothesized that varying landscape configuration across scenarios would impact performance of the four bundles in contrasted ways; mostly at low rather than high restoration targets due to high structural connectivity attained at high restoration targets (Gardner *et al*., 1992).

## 1. Methods

### 1.1. Virtual landscape simulations

Virtual landscapes simulations, ES models and data analyses were run using Python 3.11.3 and R 4.3.0. A list of used packages is provided in the supplementary material. The different steps and parameters of the simulations are shown in Figure 1a.

**Figure 1.**
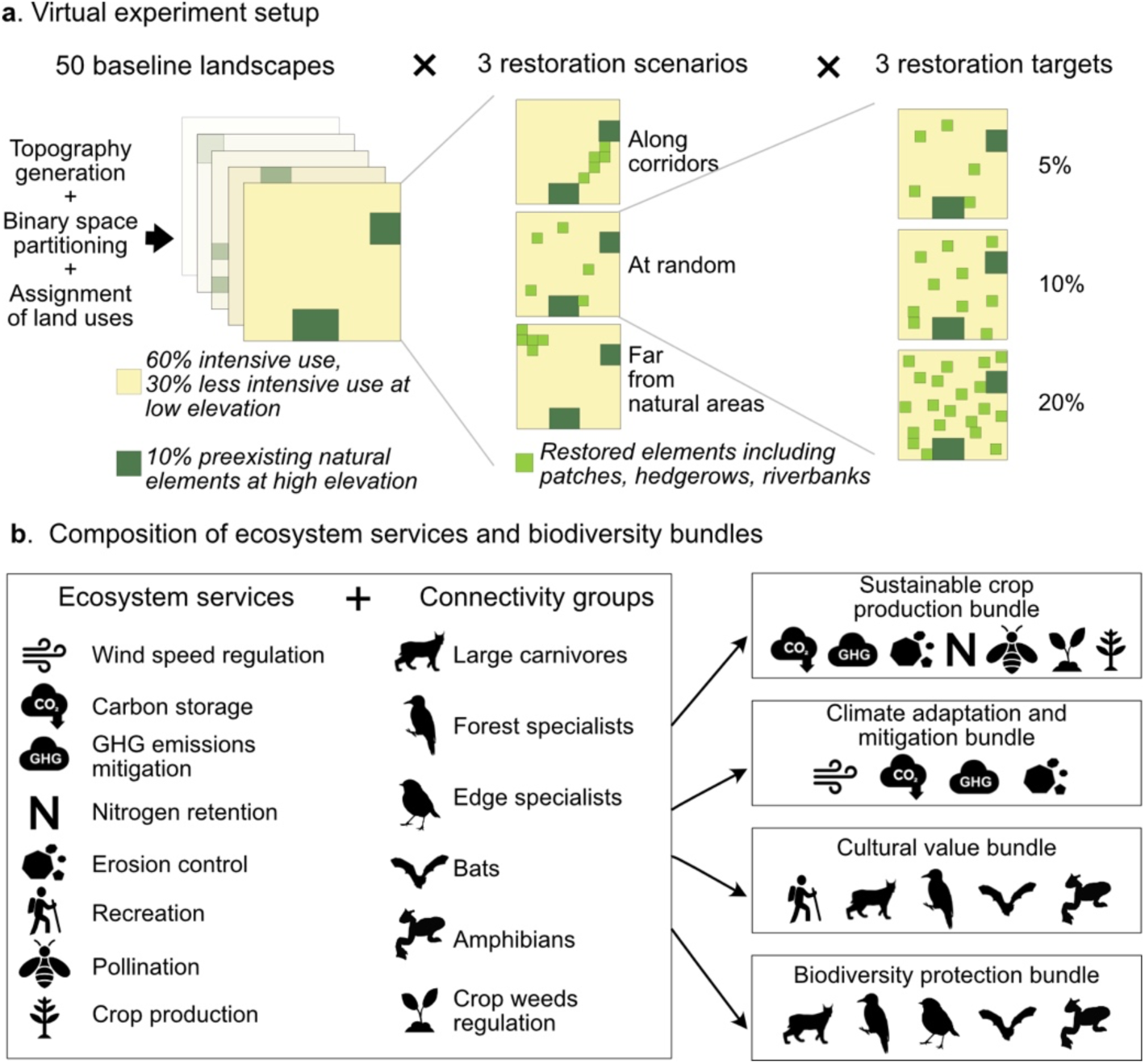
Overview of the methods. a. Combination of parameters used in the virtual experiment. For conciseness, only three land use classes are shown (preexisting natural areas, other land uses, newly restored areas); in reality baseline landscapes included six land use classes (see methods and supplementary material). b. List of ecosystem services and connectivity groups models and composition of the bundles.

#### 1.1.1. Baseline landscapes

Fifty baseline landscapes were created following the procedure presented in Richards *et al*., 2024 (see details in the supplementary methods). Briefly, each landscape comprised 2000 × 2000 cells with a 25 m grain and therefore a 50 × 50 km extent. The topography was initialized using randomly directed slope added to Perlin noise to produce hilly landscapes (Etherington, 2022). Each landscape was divided into 2000 rectilinear parcels using binary space partitioning (Etherington *et al*., 2022), with higher partitioning probability (resulting in smaller parcels) at low elevations. We then applied a baseline landscape composition reflecting an intensive agricultural landscape, with 60% intensive agricultural use, 10% natural cover, and 30% less intensive use. Human-dominated landcovers (crops, intensive grasslands, planted forests) were first processed sequentially. For each land cover, a random parcel among the smallest available was selected. The land cover was then expanded to the nearest parcel from the smallest ones available, until each land cover class had reached its specified maximum proportion of the landscape: crops 6%, intensive grass 54%, and forest plantations 10%. Then, extensive grassland was assigned to the 20% flattest parts of the remaining landscape, natural forest was assigned to the 5% steepest parts, and shrubland was assigned to the remaining 5%. We focused on landscapes with high anthropogenic footprint because their restoration is critical both in terms of biodiversity and ecosystem services (Grass *et al*., 2019); they are also representative of landscapes throughout agricultural temperate areas: for instance, the Bièvre plain in South-East France is composed of 70% intensive arable land (both grasslands and croplands) and 25% less intensive land uses (extensive grasslands, forests among which mostly production forests) with few seminatural elements (Vannier, Bierry, *et al*., 2019; Vannier, Lasseur, *et al*., 2019). At larger scale, 67% of 1km^2^ landscapes throughout Europe were classified as intensive or very intensive by van der Zanden *et al*., (2016) based on their dominant land use (intensive cropland and/or grassland) and structure.

#### 1.1.2. Scenarios of connectivity restoration

We conducted restoration experiments by converting human-dominated land covers into semi-natural land covers (natural forest patches, riparian forest corridors, or hedges). We generated nine restoration scenarios for each landscape, by factorial combination of three spatial restoration scenarios and three restoration targets (area restored representing 5, 10 or 20% of the landscape; Figure 1a). The largest restoration area was chosen to reach the proposed target of 30% of natural woody vegetation (Arroyo-Rodríguez *et al*., 2020) or 30% restored area from the 30×30 target (CBD, 2022), with smaller restoration targets representing half and a quarter of this value. The three spatial restoration scenarios stylise different restoration options that generate contrasted configurations of natural elements in a landscape, generating variation in various landscape metrics (e.g. aggregation level, patch size, distance between patches and edge length; Figure 2). As these metrics were often strongly related we only present results for aggregation.

**Figure 2.**
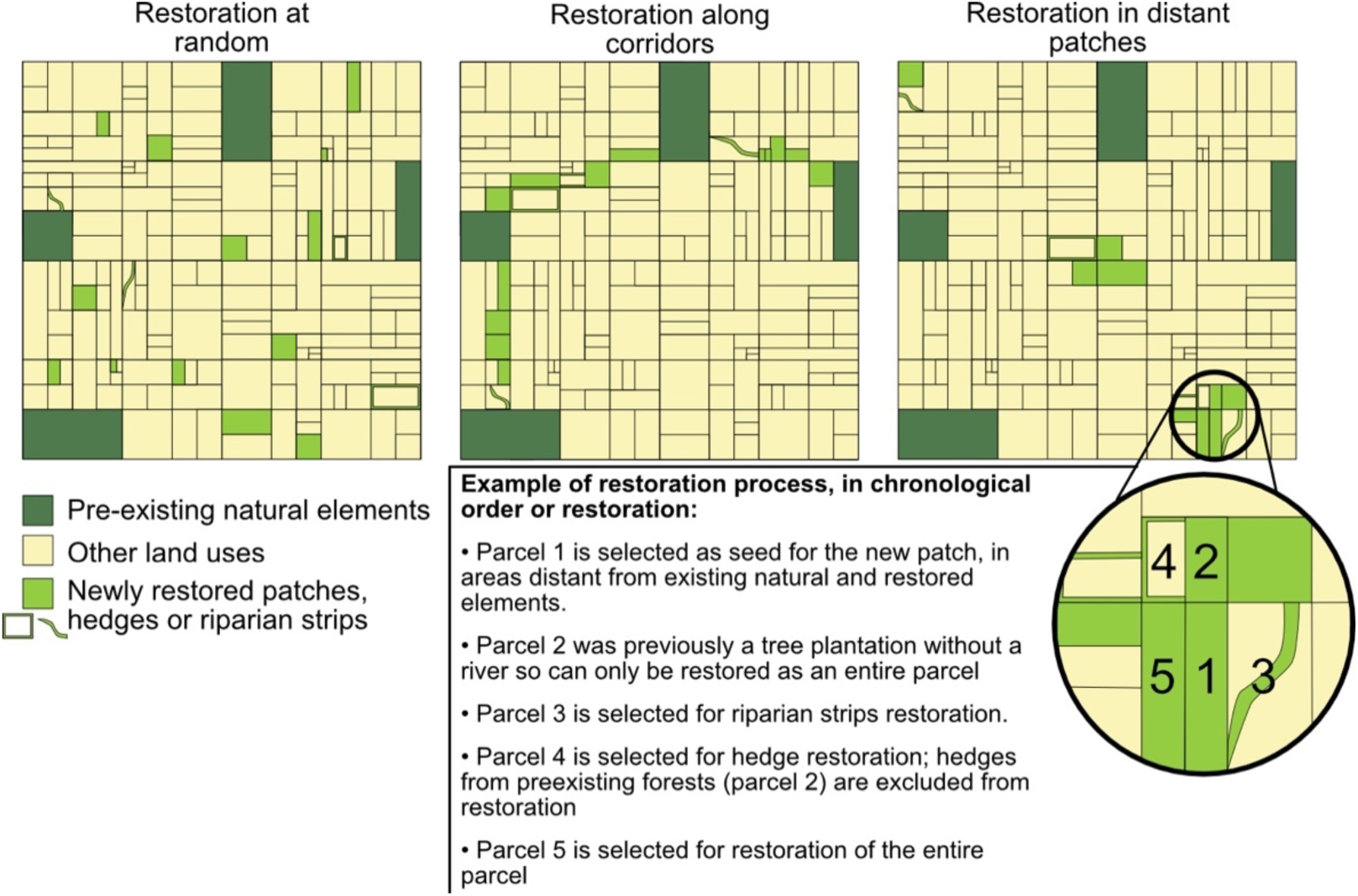
Example of restoration scenarios outcome and process at 10% restoration target. For conciseness, only three land use classes are shown: preexisting natural areas, other land uses, and newly restored areas. Outcomes on full-size landscapes showing all land uses shown in Figure S1.

Restoration at random spatial locations provided a null model where restoration had no spatial planning and was expected to generate low aggregation, because (under a certain restoration threshold) selected parcels are as likely to be isolated rather than next to an existing or restored natural parcel. Restoration in patches most distant from existing natural elements aimed to improve the ecological value of the areas in the landscape most devoid of natural elements. While potentially partly isolated from existing natural areas, these patches could still contribute to landscape connectivity for long-dispersing species and provide ecosystem services to their surroundings. This scenario was expected to generate high aggregation. Finally, restoration along potential corridors (here defined as least-cost paths, irrespective of the underlying land use, between natural areas) represented the largest-scale spatial planning process, in which restored parcels are selected along potential ecological corridors; generating intermediate aggregation. This intends to model the green infrastructure approach, defined as “strategically planned network of high-quality natural and semi-natural areas, designed and managed to provide the greatest amount of ES and protect biodiversity, both in rural and urban settlements” (European Commission 2013) which largely relies on core areas connected by ecological corridors (Ortega *et al*., 2023). Details of the simulation process for each scenario and examples of resulting landscapes can be found in the supplementary material.

At each restoration step, one of three restoration options was randomly selected: the whole parcel, hedges, or riparian strips; thus all final landscapes contained a mix of these three elements. In the case of restoration of the whole parcel, one parcel was selected among all parcels available for restoration at this step (as determined by the restoration scenario) and entirely converted to natural forest (Figure 2, parcel 1). When hedge restoration was selected, one parcel of crop or intensive grassland was selected and pixels adjacent to its edges restored. Only edges that were not adjacent to existing hedges or forests were restored (Figure 2, parcel 4). When riparian strip restoration was selected, one parcel crossed by a river and among the parcels available for restoration was selected, and pixels within the parcel directly adjacent to the river were converted (Figure 2, parcel 3). If one restoration option was not available (e.g. no more parcels with rivers) another option was randomly selected (Figure 2, parcel 2). All pixels converted to forests, hedges or riparian forests counted towards the restoration target. In terms of biophysical and ecological characteristics for ES assessment (see below), hedges were considered mostly similar to shrubs, and riparian vegetation to forests, based on their structural characteristics.

### 1.2. Landscape-level ecosystem services

We modelled eight ES commonly demanded from rural landscapes (Figure 1b): crop production, carbon stocks, greenhouse gas emissions mitigation, pollination, erosion mitigation, nutrient retention, wind protection and landscape recreation value. Most ES were modelled following published methods (Lavorel *et al*., 2022; Richards *et al*., 2024); wind protection modelling followed Vigiak *et al*. (2003). Table 1 provides an overview of the models used, with details available in the supplementary methods and Table S1. All indicators represent positive impacts for society.

**Table 1.** Description of the ecosystem service models – see supplementary methods and Table S1 for details of each model.

| Ecosystem service | Pixel-level ES | Landscape-level ES |
| --- | --- | --- |
| **Greenhouse gas emissions** | Look-up table; data for each land cover from Thomas *et al.*, (2020) | Sum of pixel-level emissions |
| **Carbon stocks** | Look-up table; data for each land cover from Mason *et al.*, (2012); Case and Ryan, (2020); Thomas *et al.*, (2021) | Sum of pixel-level stocks |
| **Nitrogen retention** | InVEST nutrient delivery ratio model (Natural Capital Project, 2023). Sources of nutrients and transport rate per land use from from Lavorel *et al.* 2022. | Ratio of nitrogen load/nitrogen export across the landscape |
| **Erosion control** | Universal Soil Loss Equation (Renard *et al.*, 1991) with vegetation parameters based on expert assessment (Lavorel *et al.* 2022) | Minus mean annual erosion rates across the landscape |
| **Pollination** | Proportion of cropland within 500m of pollinator habitat and thus assumed to receive a pollination service (Schulp, Lautenbach and Verburg, 2014) |  |
| **Recreation** | Recreation opportunity spectrum includign three components: proximity to water, elevation, and attractiveness of the patch (Lavorel *et al.*, 2020) |  |
| **Meteorological shelter: Wind protection** | Friction velocity reduction factor F calculated from the height, width and porosity of all woody elements (Vigiak *et al.*, 2003) | minus the average of F across non-woody areas |
| **Crop production** | Look-up table; data on yield extracted from (Agreste 2008-2012) and (Département de l'Isère and Direction Départementale des Territoires, 2019) | Sum of production across the landscape |

### 1.3. Functional connectivity for biodiversity

#### 1.3.1. Definition of connectivity groups

In order to assess the impact of restoration scenarios on landscape connectivity for a range of species with distinct characteristics, we adopted a generic focal species (GFS) approach. GFS is a conceptual species, whose profile consists of a set of ecological requirements or traits which reflect the likely needs of real species (Watts *et al*., 2010). For simplicity, in the following we call GFS “connectivity groups”. We modelled six connectivity groups (Figure 1b), differing in their habitat preference, dispersal distance (short / long), and habitat size requirement (small / large, Table 2, supplementary methods). We fixed the short and long dispersal distances to 1km and 20km respectively, to cover a range of dispersal distances around the reference dispersal distance of 10km used in previous studies (Saura *et al*., 2017), although the whole range of dispersal distances (up to 100+km) was not covered due to the relatively small extent of our landscapes. The minimal patch sizes to 0.5km^2^ and 10 km^2^ were chosen based on plausible body mass for different connectivity groups (Prima et al. in review), transformed into minimum habitat size based on Tucker, Ord and Rogers (2014). While these generic parameters might not correspond to any specific species, we provide one illustration of a real species that could broadly correspond to each group (Table 2). Some species with dispersal distance outside the chosen range of dispersal distances and patch size (some small rodents and amphibians with small dispersal distance, or large vertebrates with long dispersal distances) might not fully be represented; so we also provide sensitivity analyses with smaller or larger dispersal distances and patch sizes. Higher connectivity was considered as positive impacts, except for invasive weeds for which it was considered negative and inversed (“invasive weeds control”).

**Table 2.**
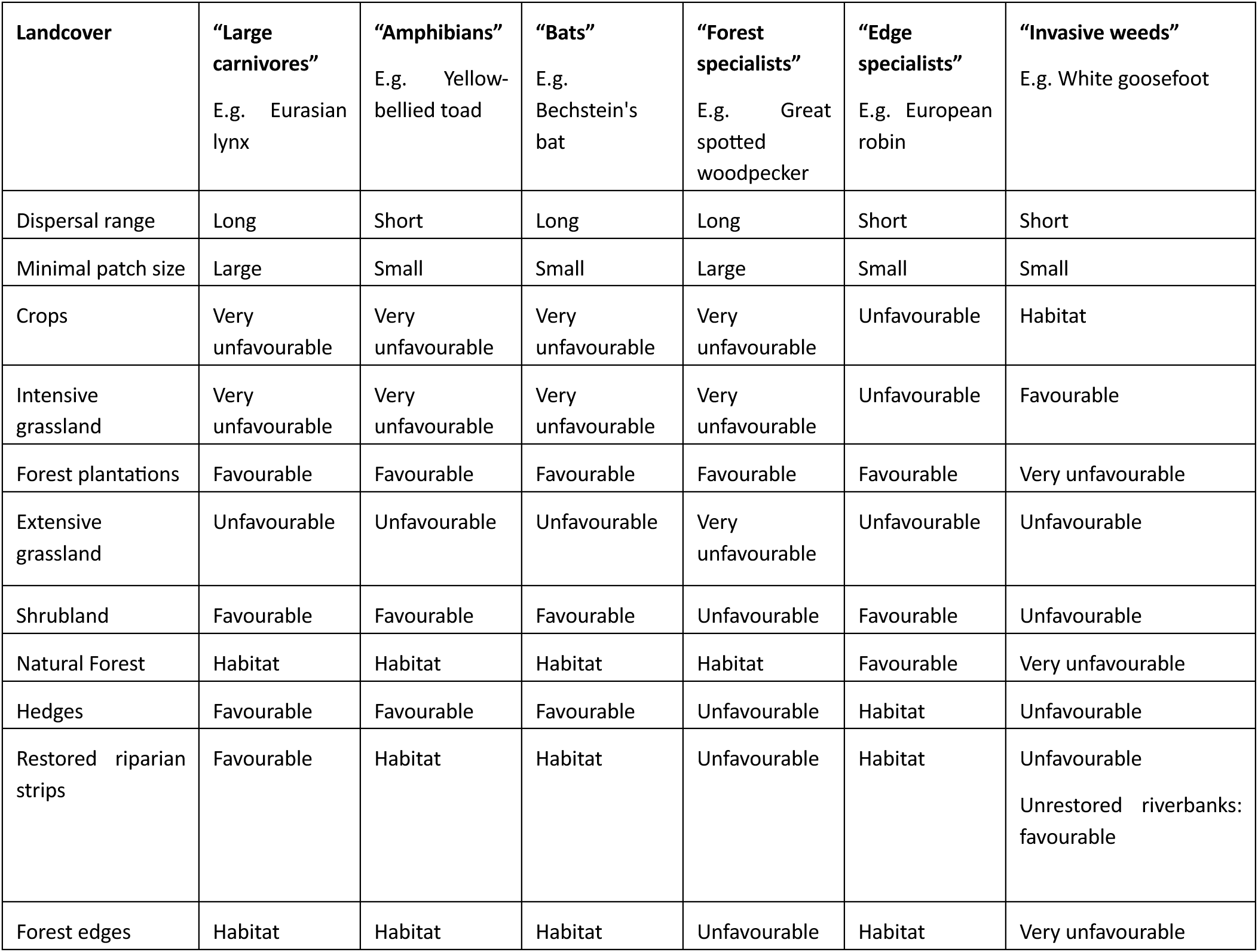
Parametrisation of connectivity groups. Each group has contrasted habitat requirements and dispersal characteristics, in order to represent the effect of landscape connectivity on a breadth of connectivity groups. Land uses are classified based on their suitability for each group: “habitat” indicates core niches with maximal suitability, in which organisms can move freely while “favourable”, “unfavourable”, “very unfavourable” describe land uses with decreasing habitat suitability (and increasing resistance to movement).

#### 1.3.2. Parametrization of matrix resistance

We set the habitat suitabilities H to 100% for habitat, 70% for favourable matrix, 30% for unfavourable matrix, and 10% for very unfavourable matrix. Land cover maps were converted to resistance (R) maps, i.e. the impermeability of each land cover for the movement of organisms following Keeley, Beier and Gagnon, (2016):

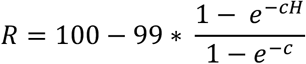

With c a transformation factor determining the shape of the curve linking habitat suitability and resistance. Because of the uncertainty of the relationship between suitability and resistance, we tested different c values; only results for c = 8 are shown in the main text.

#### 1.3.3. Landscape connectivity assessment

For each group, we estimated functional connectivity by defining dispersion kernels (Figure S2). Source patches were identified as contiguous areas of habitat that were at least as large as the group’s minimal patch size. We attributed a resistance value of 1 to habitat pixels, except those located in source patches which had a resistance of 0. While a null resistance only exists in theory, we considered it negligible compared to the resistance of other land uses, and thus unlikely to affect the connectivity between habitat patches; within-patch connectivity was not considered. This simplification allowed to considerably decrease computation time by selecting only one source pixel per source patch, with the dispersal costs being counted only from the edges of the source patch.

Then, we calculated dispersal kernels as the area reachable from each source patch, conditioned by the dispersal distance through least-cost paths. Each pixel contributed to the total distance proportionately to its resistance, i.e. a species with a dispersal distance of 1km could move 1km through (non-source) habitat, or 100 m through land use of resistance 10. The landscape-level connectivity indicator for each group was then defined as the probability that two randomly sampled points in the landscape belonged to the same resistance kernel, calculated across 10^6^ points (equivalent of 1 - the landscape-scale *Division* metric, Jaeger, 2000). This index thus increases with the proportion of the area covered by the dispersal kernels, and decreases with the number of kernels. If the calculated index was equal to 0, we set it to the theoretical minimum of 10^-6^ instead, to allow for further data transformation.

### 1.4. Data analysis

We described the composition and configuration of each baseline and restored landscape as the landscape-level proportion of each land use class and the aggregation index of all natural elements combined (forests, shrubland, hedges and restored riparian strips). Landscape aggregation was calculated as the number of like adjacencies divided by the theoretical maximum possible number of like adjacencies for natural elements for the entire landscape (He, DeZonia and Mladenoff, 2000).

ES and connectivity groups were bundled into four landscape management objectives reported in studies focusing on how people’s priorities for different ES are often associated (“demand” bundles, hereafter “ecosystem services and biodiversity bundles” or “bundles” - Figure 1b) (Brown, Helene Hausner and Lægreid, 2015; Ament *et al*., 2017; Lavorel *et al*., 2019; Zoderer *et al*., 2019; Peter *et al*., 2022): 1. sustainable crop production: crop production, invasive weeds control, carbon storage, greenhouse gas emissions mitigation, erosion control, nitrogen retention and pollination; 2. climate adaptation and mitigation: wind speed regulation, carbon storage, greenhouse gas emissions mitigation and erosion control; 3. landscape cultural value: recreation, predators, bats, amphibians and forest specialists; and 4. biodiversity protection: large predators, bats, amphibians, bird forest specialists and edge specialists. Because the distributions of the indicators were not Gaussian, each indicator was first scored from 1 to 10 based on the 10% quantile each value belonged to. When 0 values represented more than 10% of all values, they were given the minimal score of 1. The value for each bundle was then calculated as the average score across indicators.

For each restored landscape, we calculated the relative change in each bundle as the difference between its value in the baseline and restored landscape, divided by the baseline value. We also calculated the relative change in the aggregation of natural elements following the same procedure.

We conducted within-group PCAs, centred on the restoration target, on i) compositional and configurational characteristics of landscapes, to identify the main axes of variations across landscapes (Figure S3) and ii) ES and functional connectivity in each bundle, to explore the multidimensional response of each bundle to landscape configuration (Figure 3). In within-group PCAs, the data is centred around the mean of defined groups – here, the restoration target – before running the PCA, allowing to investigate the association between the variables irrespective of the group.

**Figure 3.**
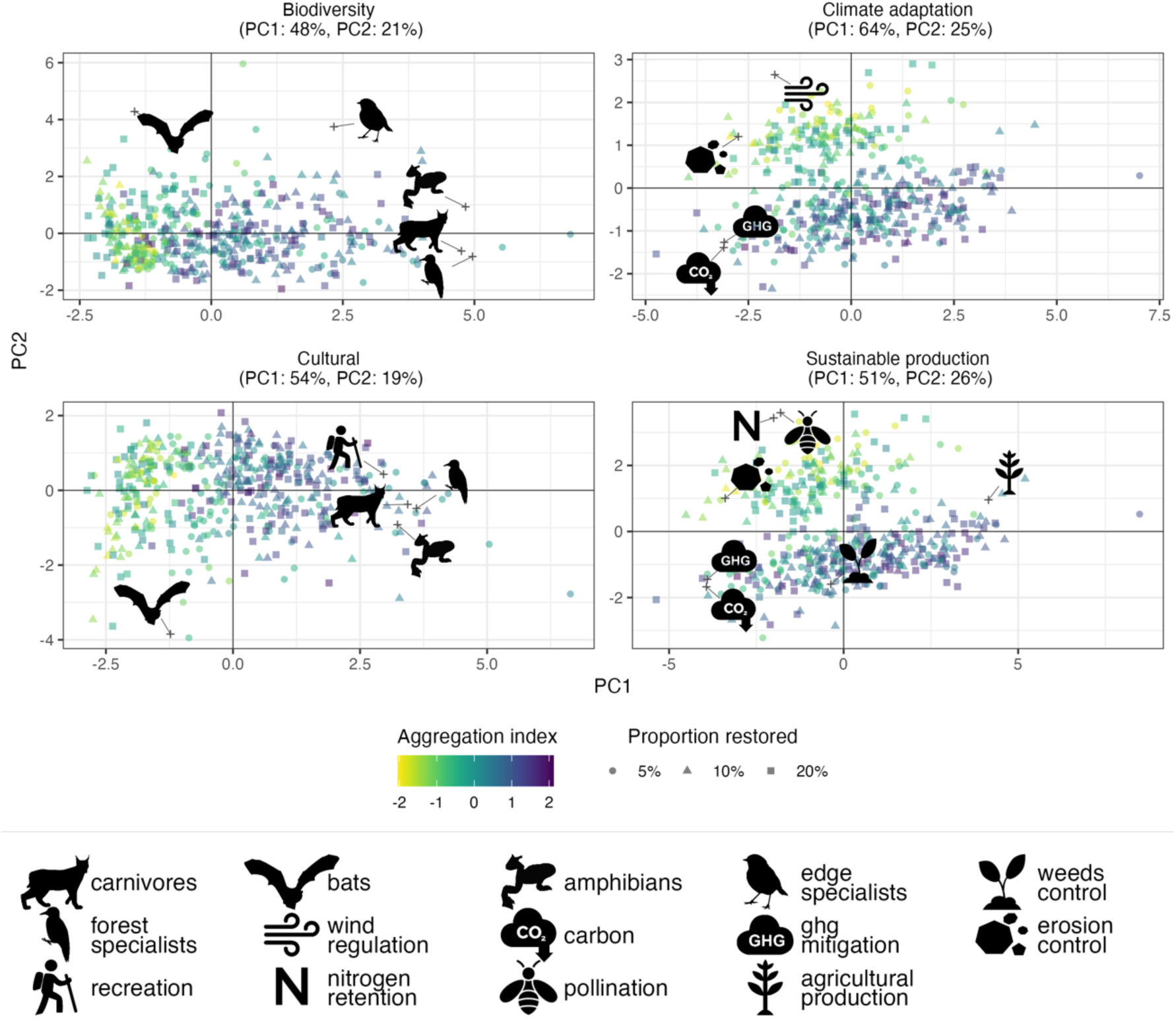
Within-group PCA of each bundle. The PCA were centred on the restoration target (i.e., the effect of the target was removed before PCA), hence the absence of pattern for this variable, shown by dot shape. The dot colour represents the aggregation of the natural areas in each landscape (yellow = low aggregation, purple = high aggregation). The crosses show the precise score of each variable. N = 150 for each panel.

To assess the difference between various restoration targets and restoration scenarios, we compared the changes in bundles across restoration targets, and across scenarios within restoration proportions, using ANOVAs. We regressed the changes in each bundle with the changes in aggregation of natural elements in the landscape in linear models, in which we compared the slopes across the different restoration targets.

## 2. Results

### Landscape composition and configuration

Hedges and riparian strips represented a small fraction of the landscapes (0.26±0.05% to 0.91±0.09% for hedges and 0.15±0.05% to 0.61±0.12% for riparian strips at 5% and 20% area restored, respectively). When excluding the effects of the area restored, the main axis of variation between landscapes (PC1: 38%; Figure S3) ranged from landscapes characterised by low aggregation and distance between patches with (comparatively) higher covers of hedges and riparian strips, to landscapes with high aggregation with higher covers of forest patches. The second axis (PC2: 20%; Figure S3) differentiated restored landscapes with comparatively high proportions of extensive grasslands, from restored landscapes with high proportions of intensive grassland. The differences in land cover proportions resulted from the different areas prioritised by the scenarios for restoration. For instance, in the random restoration scenario, a higher proportion of the total restoration target consisted of hedges and riparian strips; this was because at each step of the restoration procedure there were more available options for these types of restoration (Figure S4).

### Ecosystem service and biodiversity bundles

Although the selection of the four bundles was based on common management objectives and not *a priori* on their spatial sensitivity, the ES and connectivity groups within each bundle were usually promoted by similar landscapes. At fixed restoration target, all ES and connectivity groups related to *biodiversity* and *cultural value* were promoted by high aggregation, with the exception of bats, which are not constrained to large patches. Conversely, most ES for *climate adaptation* were either independent from aggregation (carbon, GHG mitigation), or promoted by low aggregation (wind regulation, erosion control). *Sustainable production* included ES promoted by both low (pollination) or high (edge specialists) aggregation (Figure 3).

### Responses of ES and biodiversity bundles to restoration scenarios

All bundles responded positively to restoration, with the strongest effects observed for *biodiversity* and *cultural value* (up to 5.8±0.7% and 5.7±0.7% change, respectively, at 20% restoration, Figure 4). Both bundles were promoted by restoration along corridors or in distant patches better than by random restoration, especially at low restoration targets. ES and connectivity groups within these bundles showed different response shapes to the restoration target. While recreation and connectivity groups with low habitat size requirements increased gradually with restoration target, groups which required large habitat patches had a threshold behaviour (Figure 4). At 5% restoration target, only the restoration scenario in distant patches provided sufficient patch sizes for these species. At 10% restoration target, restoration along corridors sometimes generated sufficient patch size, but random restoration almost never. At 20% restoration target, minimal patch size was reached in all scenarios.

**Figure 4.**
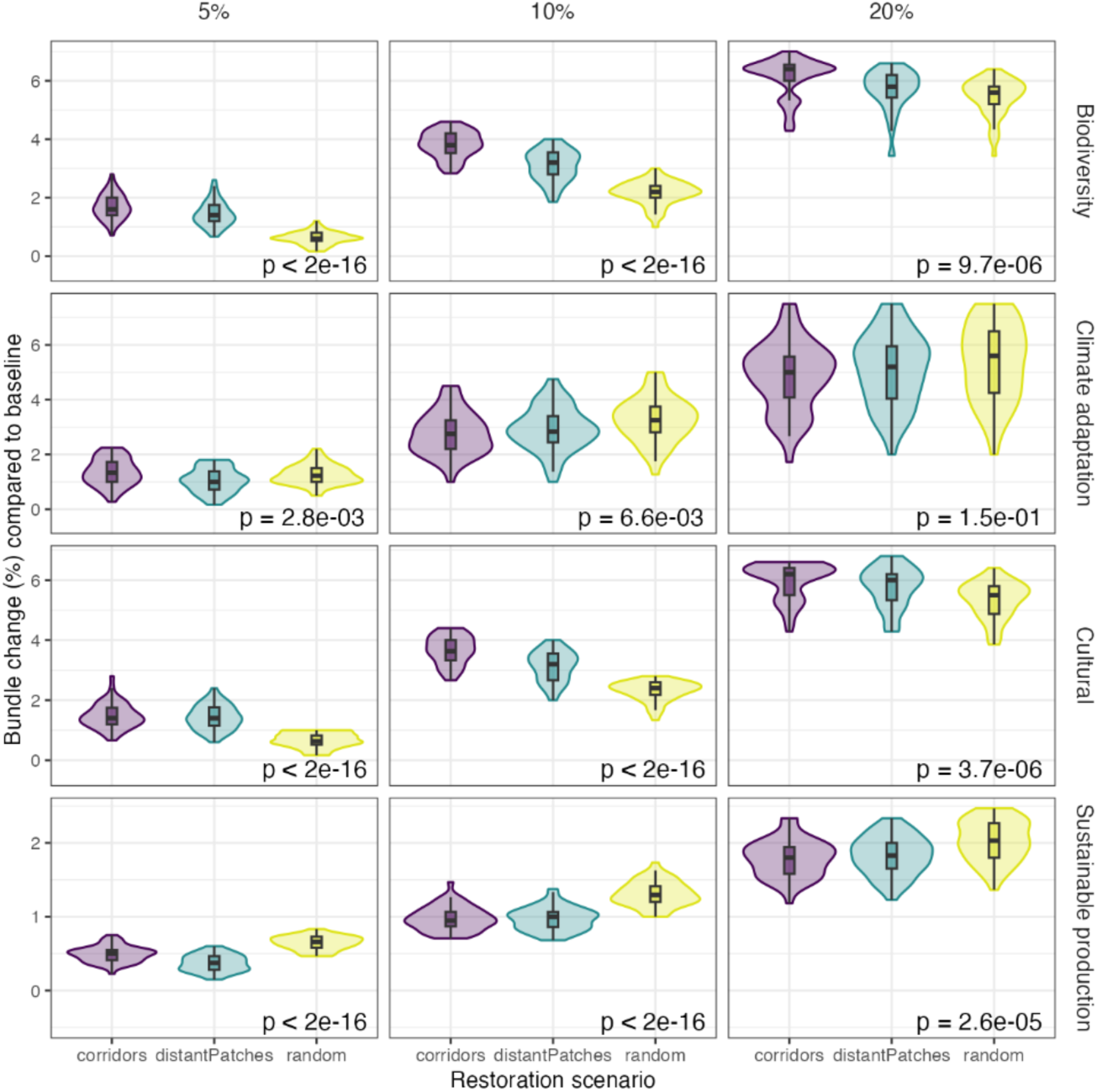
Change in bundle values per restoration scenario for each restoration target. Violin plots show the distribution of the data points, box plots show median and interquartile range. P-values were extracted from anovas run for each bundle and each restoration target independently, with fdr correction for multiple testing. N = 150 for each panel.

ES related to *sustainable production* and *climate adaptation* also increased with restoration, except agricultural production decreasing due to conversion of agricultural land (Figure 4). This trade-off meant that overall *sustainable production* only increased up to 1.9±0.3% at 20% restoration. The responses of *sustainable production* and *climate adaptation* were also modulated by the restoration scenarios, as randomly located restoration yielded higher values for both bundles, especially at low restoration targets (Figure 4). This was mainly due to the strong positive response of specific ES to this scenario, like pollination, wind speed regulation and nitrogen retention; although other ES such as carbon storage or GHG mitigation also showed weaker but significant responses to the restoration scenarios (Figure 4).

### Interactions between configuration and restoration target

The contrasted effects of the restoration scenarios on bundles corresponded to changes in the configuration of natural elements in the restored compared to baseline landscapes (Figure 5). Strong increase in aggregation promoted *biodiversity* and *cultural value* (p < 10^-6^) because many species require habitat patches of sufficient area. The strength of these effects varied with the restoration target (interactions aggregation: restoration target p < 10^-6^ for *biodiversity* and p = 0.01 for *cultural value*), with weaker but still significant slopes at high restoration target (Figure 5). On the contrary, *sustainable production* and *climate adaptation* decreased with increasing aggregation (restoration target: p < 10^-6^, aggregation p < 10^-6^; weak interaction: 0.05 < p ≤ 0.1, Figure 5). This is because their component ES mostly rely on certain habitats (e.g. pollination) or physical features (e.g. windbreak) providing ES to their surrounding (mostly) independently from their own areas, resulting in larger gains for small, interspersed natural elements in the landscape. Some ES also displayed no response to aggregation although they varied across scenarios (carbon, GHG emissions, Figure S7).

**Figure 5.**
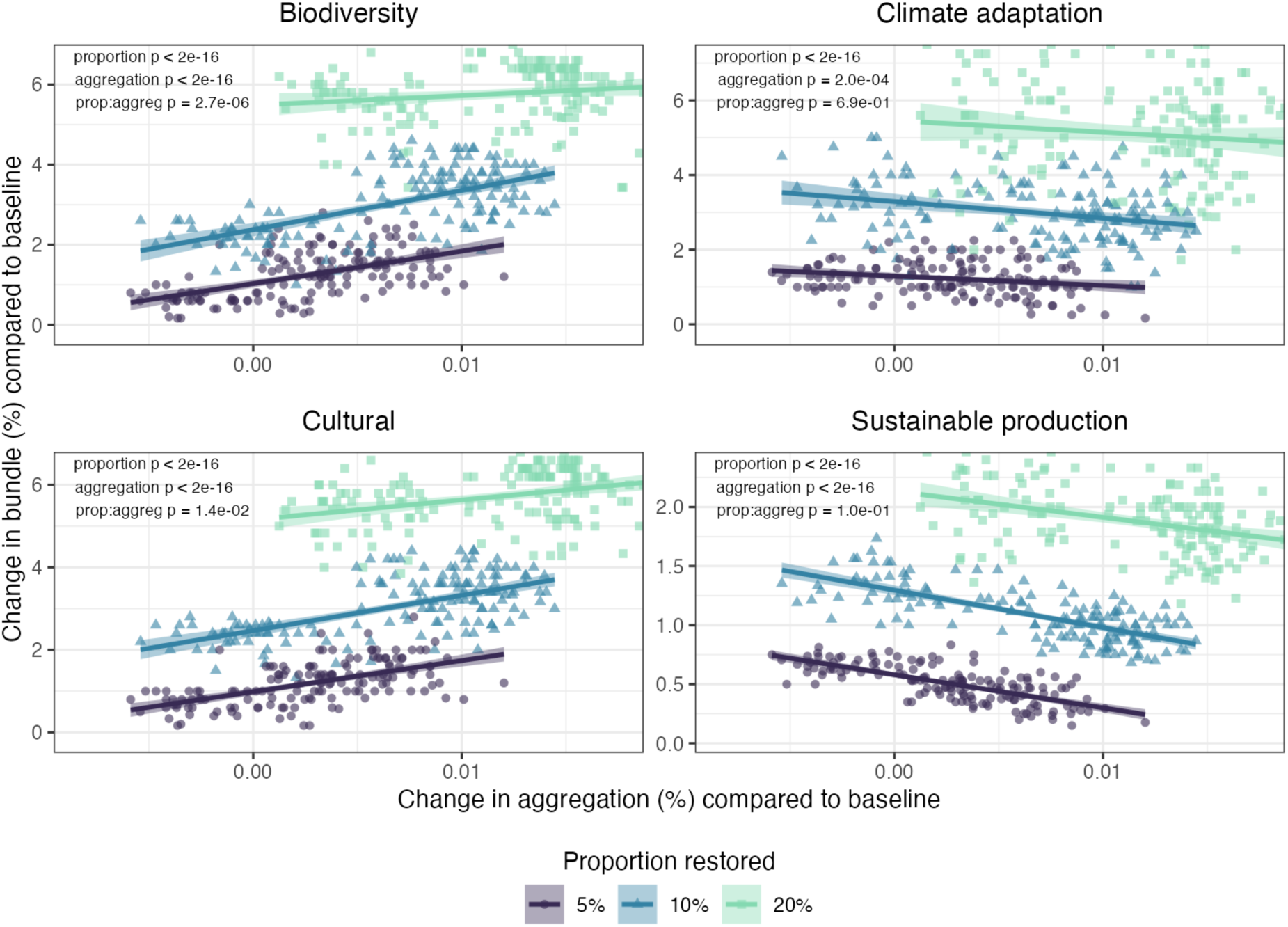
Effect of aggregation of natural elements in a landscape on each bundle, depending on the total proportion of the landscape restored. Model results are shown in Table S2. N = 150 for each colour in each panel.

### Sensitivity analyses

Larger conversion factors (Fig. S 9-S 12), dispersal distances (Fig. S 17-S 20), smaller patch sizes (Fig. S 13-S 16), and smaller target sizes for restoration of distant patches (Fig. S 21-S 24) decreased the sensitivity of the biodiversity and cultural bundles to restoration scenario and landscape aggregation, but did not change the results qualitatively.

## 3. Discussion

Using virtual landscape experiments, we tested the impact of contrasted restoration scenarios on ES and connectivity groups within production landscapes. We confirmed that landscape composition, here determined by the restoration target, was the main driver of all considered bundles (Duarte *et al*., 2018) whether restoration happened as large patches, along corridors, or randomly throughout the landscape. This shows that besides protecting large natural areas, other effective area-based conservation measures such as those presented here – improved hedge network or small woodland patches - could also promote biodiversity and ES within production landscapes (Dudley *et al*., 2018). Restoration scenarios mattered most at low restoration levels, consistently with previously described mechanisms of structural connectivity at 30-40% of natural elements in a landscape (Gardner *et al*., 1992), corresponding to our maximal restoration target. However, for restoration targets lower than this threshold, different bundles were promoted by different landscape configurations, and thus the specific restoration strategies influenced the outcome of common management objectives.

### Bundle responses to landscape configuration

*Biodiversity* was promoted by scenarios generating high aggregation of natural elements. There has been much debate regarding the effects of landscape fragmentation on biodiversity: on the one side, small patch size increases possibly negative edge effects, and might reduce population sizes below viable levels (Levins, 1970), especially in larger-sized organisms (Uezu, Metzger and Vielliard, 2005). On the other hand, fragmented landscapes have been shown to generally host higher biodiversity (Fahrig, 2017). Here, rather than the number of species, we focused on landscape connectivity, which promotes gene flow, population mixture, and overall long-term resilience, for instance by allowing species to track their climatic niches (McGuire *et al*., 2016). Low aggregation of natural habitats had negative effects on most connectivity groups, either because of the lack of sufficiently large habitat patches (e.g. for large predators) or due to the inability of short-dispersal species to move between isolated patches. This was so even though the minimal patch size for high habitat requirements (10km^2^), chosen to represent a large range of species, was substantially lower than actual patch sizes reported for large predators (e.g., unfragmented habitat of 25-30km^2^ needed for lynx, Farrell *et al*., 2018). The exception to this was organisms with small patch requirements and large dispersal distance, consistent with highly mobile organisms being considered to be generally more resilient to habitat fragmentation (Ewers and Didham, 2006).

Some of the ES included in *climate adaptation* and *sustainable production* such as greenhouse gas emissions were not considered to primarily depend on patch area, and thus modelled as simple lookup tables without spatial mechanisms, resulting in insensitivity to landscape configuration. This could be improved in future studies, as landscape configuration has been shown to also impact these ES either directly (Didham *et al*., 2015) or indirectly through effects on supporting soil biota (Le Provost *et al*., 2022). Still, these ES varied across restoration scenarios, mostly due to changes in land cover proportions as the scenarios prioritised different areas of the landscape, and thus different land covers, for restoration (Richards *et al*., 2024). Conversely, many ES require interspersion of natural and human habitats (Mitchell *et al*., 2015). At fixed restored area, lower aggregation with small patches dispersed throughout the landscape decreased the average distance between crop areas to the nearest natural habitat, which promotes high ES flow (Mitchell, Bennett and Gonzalez, 2015): for instance, wind speed regulation depends more on distance to woody elements than on their size; thus open areas benefit from many small elements more than from few large ones. Besides, natural elements such as riparian strips or hedges promote water infiltration and nutrient retention in soil and roots (Thomas and Abbott, 2018). Their efficiency depends primarily on their length, rather than width (Brumberg *et al*., 2021), encouraging the restoration of long, thin strips of (semi-) natural vegetation.

### Virtual landscape approach

Virtual landscape experiments have been used to assess landscape pattern effects on biota depending on their habitat requirements and dispersal abilities (Simpkins *et al*., 2018). Recent studies have used virtual landscapes experiments to identify general rules regarding the impact of landscape composition, heterogeneity (van der Plas *et al*., 2019) or configuration (Mitchell, Bennett and Gonzalez, 2015; Richards *et al*., 2018; Lavorel *et al*., 2022) on ES in a range of simple, controlled scenarios. Here, we go beyond previous studies focusing on restoration scenarios on ES supply and area of suitable habitat (Richards *et al*., 2024) by expanding the analysis to integrate quantitative tolerance for different land uses of multiple connectivity groups. This demonstrates the applicability of landscape experiments to jointly assess the impact of landscape connectivity restoration on multiple ES and biota. Large-scale, remote sensing-based assessment of real-world landscapes (Oehri *et al*., 2020), selected to display independent variation of composition and configuration could help validate the general principles derived from such virtual landscape experiments.

We focused on a small set of restoration scenarios options, targeting simultaneously linear elements (hedgerows and riparian strips) and forest patches of varying size, which support different ES and biota (Valdés *et al*., 2020; Ortega *et al*., 2023). We also focused on restoration involving the planting or natural regeneration of trees, which might not be appropriate in all ecosystems (Veldman *et al*., 2019). Further exploration of restoration options could investigate the relative benefits of restoring linear elements or patches only, as well as more complex restoration options involving other types of ecosystems, such as the restoration of semi-natural pastures and grasslands (Richards *et al*., 2024), as they host distinct biota and ES (Bardgett *et al*., 2021; Lyons *et al*., 2023) and are critical to biodiversity conservation and functionality or working landscapes (Smith *et al*., 2013). Besides, the implementation of conservation measures that improve the quality of the matrix (e.g. flower strips, scattered trees) would decrease the contrast between natural and human-dominated land covers, overall improving the permeability for wildlife and ES supply (Arroyo-Rodríguez *et al*., 2020). Finally, a key step towards understanding the effect of restoration strategies on ES and biodiversity is to integrate human elements such as roads or settlements in landscape modelling (Etherington et al. 2023).

### Application to real landscapes

Simulated scenarios captured a stylised, yet representative spectrum of spatial configurations for restoration demonstrating their critical impact on outcomes for biodiversity, multiple ecosystem services and their beneficiaries. Their application to real-life landscapes requires taking their biophysical, ecological and social context into account. First, landscape composition and configuration are not snapshots in time, but reflect their history of land use and management which will influence where restoration is ecologically relevant and socially acceptable, as well as the outcome of restoration actions (Sandberg, 2016). For instance, past land use affects biodiversity and the ES it underpins (Bommarco *et al*., 2014; Martin *et al*., 2020). Implementation of the restoration scenarios is also constrained by social factors. While random-like restoration relies on multiple landowners independently initiating small-scale restoration actions, restoration along corridors or in large isolated patches require collaboration and large-scale spatial planning. Such coordination requires collective action processes and aligning motivations (Faure, Mouysset and Gaba, 2023; Polyakov *et al*., 2023). Here, we showed contrasting impacts of landscape restoration scenarios on different ecosystem bundles, reflecting varying outcomes across the diversity of stakeholder motivations, values and demands. Stakeholders often prioritise, at least to some extent, several ES bundles; and the relative priority given to each influences which landscapes would best fit their needs (Neyret et al., 2023). Different configurations could thus favour the ES and connectivity groups most valued by different stakeholders (Boesing *et al*., 2024), highlighting the difficulty to identify restoration options that fit a wide range of biodiversity conservation, sustainability and development goals. This also suggests that combining diverse restoration strategies at landscape or regional level is likely needed, for instance with areas containing large restored patches for biodiversity, and other finer green infrastructure elements to promote other ES in working landscapes. Similar designs have been suggested when advocating the combination of large and small forest patches, embedded in a high-quality landscape, to conserve biodiversity (Grass *et al*., 2019; Arroyo-Rodríguez *et al*., 2020), and examining the required combination of landscape-level interventions for transitioning to multifunctional agricultural landscapes (Garibaldi *et al*., 2023). Participatory processes involving diverse land users and managers could support the identification of options that are tailored to local contexts and mitigate trade-offs (Barnaud *et al*., 2023).

## Conclusion

Using virtual landscape experiments, we explored the effects of different restoration scenarios on four ES bundles, representing different values associated to productive landscapes. We showed that besides the restoration target, the configuration of restored elements has a large impact on the outcomes of restoration. Trade-offs appear mostly at low restoration targets, were the bundles had contrasting responses to the scenarios and resulting landscape aggregation. Conversely, at high restoration levels, the effects of scenarios and configuration were weaker, meaning that landscapes provided high values for most ES and connectivity groups with little impact of the location and size of restoration elements. Our findings can facilitate the shift from isolated, site-specific restoration measures to the implementation of coordinated restoration initiatives at landscape and regional scales. Effective spatial planning for restoration needs to account for the configuration trade-offs highlighted above, as well as specific constraints related to the local context, when prioritizing areas to restore or protect; and this choice is all the more important as the overall capacity for restoration is limited.

## 1. Supplementary Methods

The analyses were run using packages RichdDEM (Barnes 2016), NumPy (Harris 2020), SciPy (Virtanen et al 2020), scikit-image (van der Walt et al 2014), NLMpy (Etherington et al. 2015) and GDAL (GDAL.OGR contributors 2020). Ecosystem services models were run in Python and R using packages terra, whitebox and rinvest. Data analyses were run in R using packages data.table, plyr, lconnect, vegan.

### 1.1. Baseline landscapes

Each landscape had dimensions of 2000 × 2000 cells with 25 m grain and therefore 50 × 50 km extent. The topography of each landscape was initialized using randomly directed slope with an elevation range of 0-100 m. Further topographic complexity was then incorporated by adding to the initial slope a Perlin noise neutral landscape model (Etherington, 2022) parameterized to produce hilly landscape features (periods = 10, octaves = 5, persistence = 0.4), then rescaled to range 100-800 m. Topographic depressions were then filled (Barnes 2016b) to calculate slope (Horn 1981) and to identify river channels based on D8 drainage (O’Callaghan and Mark 1984) that drained at least 3.75 km^2^. This process was repeated to create 50 different topographies.

In each of the 50 topographies, we applied a baseline landscape composition reflecting an intensive agricultural landscape, with 60 % intensive agricultural use, 10 % natural cover (natural forest and shrubland), and 30 % less intensive use (extensive grassland and forest plantation). Landcover distributions for each topography followed Richards et al. in prep. Each landscape was divided into 2000 rectilinear parcels using binary space partitioning (Etherington et al., 2022) where the probability of partitioning was based on elevation, resulting in higher partitioning probability, and thus smaller parcels, at low elevations.

Human-dominated landcovers (crops, intensive grasslands, planted forests) were then processed sequentially based on binary space partitioning depth. For each landcover a random parcel within the deepest set of available parcels (those which underwent most partitioning events, i.e. smallest in size and lowest in elevation), was selected and classified as that landcover. The landcover was then expanded by selecting the nearest parcel available from those parcels remaining at the lowest binary space partitioning depth. This process continued until each landcover class had reached its specified maximum proportion of the landscape: crops/horticulture 6 %, intensive grass 54 %, and forest plantations 10 %. The (semi-)natural landcovers of extensive grass, natural shrubland, and natural forest were assigned to the remainder of the landscape based on a gradient formed by the slope of the topography. Extensive grassland was assigned to 20 % of the flattest parts of the landscape, natural forest was assigned to 5 % of the steepest parts of the landscape, and shrubland was assigned to the remaining 5 % of the landscape that had intermediate slopes.

### 1.2. Reconnection scenarios simulation

We focus on three types of woody elements (hedges, riverbanks, whole parcels) and three possible configurations (random, depending distant from existing natural areas, or along potential corridors between natural areas. Only croplands and intensive grasslands can gain hedges. The outcome of the restoration scenarios on one example landscape are shown in Figure S 1.

In the random restoration scenario, we randomly selected human-dominated parcels for restoration, until the target restoration area was reached. In the scenarios where restoration targeted most distant patches, a “seed” parcel was selected in the 10 % parcels most distant from existing natural elements; then the restored patch was grown by restoring parcels adjacent to the seed until a target patch area of 10km^2^ was reached (5 and 20 km^2^ patch size targets are shown in the supplementary material). The process was then repeated with another seed until the restoration target was reached. Finally, for the restoration scenario along corridors we first identified corridors as least-cost paths between existing natural and semi-natural areas. These least-cost paths were independent on the connectivity group considered (i.e., structural rather than functional connectivity) and only relied a binary distinction between natural and human-dominated land covers (see supplementary methods). We then restored parcels along these paths; if all paths were restored before the restoration target was reached, parcels adjacent to already-restored parcels were then selected.

**Figure S 1.**
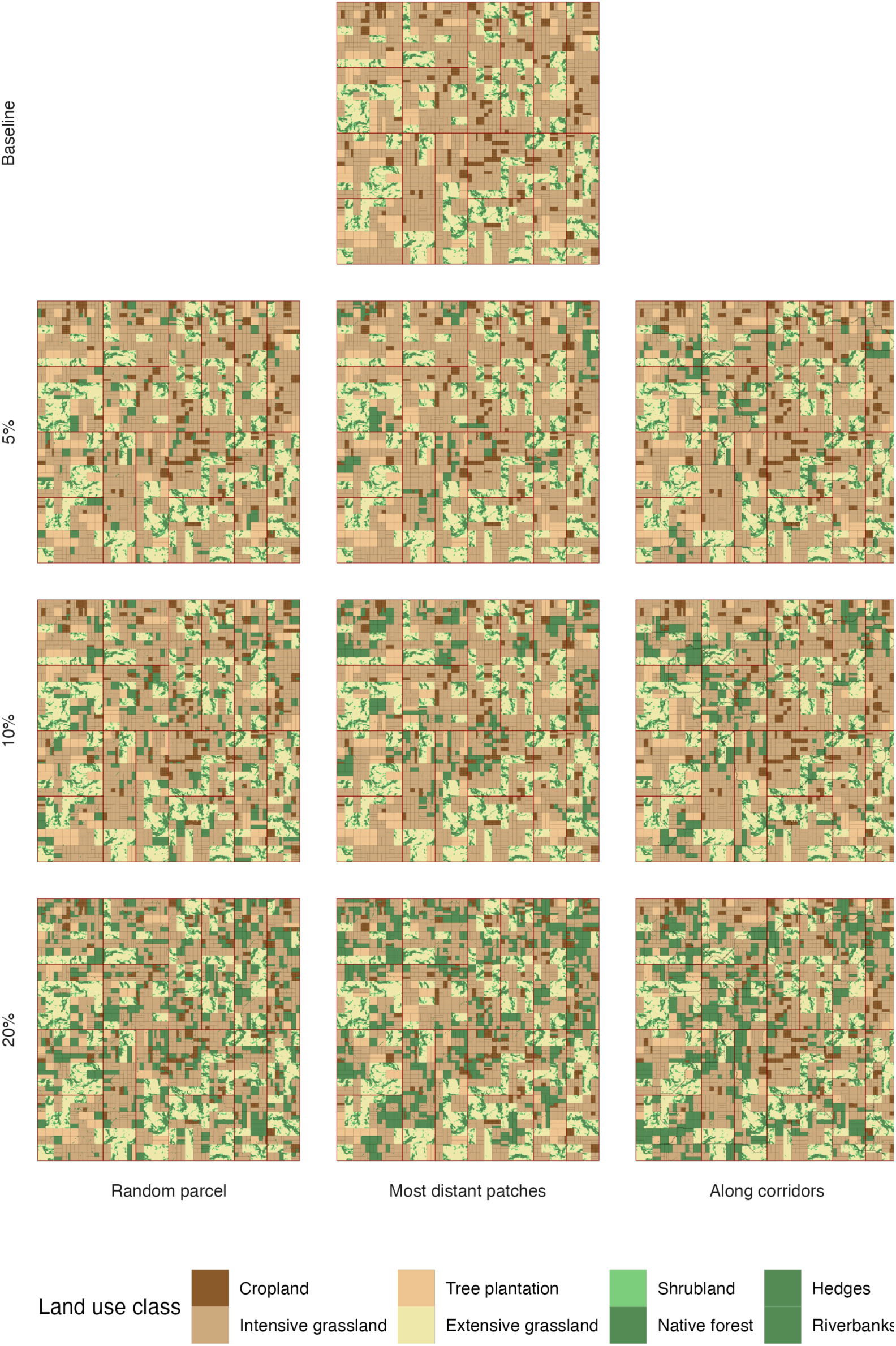
Example of restoration outcomes for one landscape

#### 1.2.1. Restoration at random

This is the most basic scenario; the same basic mechanisms are applied in the other scenarios. At each step:

- Randomly decide to restore a whole parcel, hedges along one parcel, or a riverbank portion within the intensive land uses.

- If we chose a riverbank: we choose one parcel with a riverbank in the pool of parcels to restore. If there are no such parcel left, we restore a hedge or whole parcel instead.
- Otherwise, we select randomly a parcel in the pools of parcels to restore, and we restore either the whole parcel or all its available hedges.
- Hedges next to restored hedges, restored parcels, or natural areas cannot be restored to avoid 50m-wide hedges or “useless” hedges along natural patches. However, a whole parcel can still be restored next to a previously restored hedge. A parcel can be restored only once (i.e. no parcel with both hedges and riverbanks restored)
- Repeat until target area is reached

#### 1.2.2. Restoration of most distant patches

At each step:

- All human parcels are ranked according to their distance to natural or restored areas. One “seed” parcel is selected within the 10% most distant. This is the starting parcel for the restored patch. Then, until the max patch size (10km^2^) is reached:

- All parcels adjacent to the patch (or the seed for round 1) are identified
- They are restored one by one at random as described in 1a
- The new adjacent parcels are identified
- When the maximum patch size is reached, another seed is selected and the process is repeated, until the target restoration area is reached.

#### 1.2.3. Restoration along corridors

We first identify least-cost paths between each pair of natural patches, along hedges or riverbanks that could potentially be restored. Then, all parcels located along the least cost paths are identified. At each step, parcels along the least cost path are selected and restored as described in 1a. If all parcels along the least cost path are restored, the next parcels are then selected adjacent to previously restored parcels.

### 1.3. Ecosystem service model details

**Table S 1.** Model parameters. Partly replicated from Lavorel et al. 2020.

| Landcover | Crop production | Carbon stocks | GHG emissions | Pollination | Erosion | Nitrogen retention |  | Wind protection |  | Recreation |
| --- | --- | --- | --- | --- | --- | --- | --- | --- | --- | --- |
| Model type | Look-up | Look-up | Look-up | Schulp et al 2014 | USLE | inVEST |  | Vigiak et al. 2003 |  | ROS Byczek et al. 2018 |
| Parameters | Energy production (MJ/ha) | Carbon stocks | GHG emissions | Pollinator habitat? | U | N load | N retention efficiency | Height | Barrier porosity | Attractiveness |
| Crops | 805 | 5 | 1000 | No | 0.5 | 45 | 0.4 | / |  | 0 |
| Intensive grassland | 388 | 10 | 8800 | No | 1 | 65 | 0.4 | / |  | 0.2 |
| Extensive grassland | 199 | 18 | 1000 | 1 | 0.01 | 10 | 0.6 | / |  | 0.7 |
| Shrubland | 0 | 30 | 5 | 1 | 0.005 | 4 | 0.8 | 5 | 0.7 | 0.6 |
| Exotic forest | 0 | 140 | 5 | 1 | 0.007 | 10 | 0.7 | 30 | 0.27 | 0.5 |
| Natural Forest | 0 | 200 | 3 | 1 | 0.005 | 2 | 0.85 | 30 | 0.22 | 1 |
| Hedges | 0 | 30 | 5 | 1 | 0.005 | 4 | 0.8 | 5 | 0.32 | 0.6 |
| Riverbanks | 0 | 200 | 3 | 1 | 0.005 | 2 | 0.85 | 30 | 0.26 | 1 |

#### 1.3.1. Greenhouse gas emissions

Greenhouse gas emissions were assessed from a look-up table relating the land cover of each pixel and its estimated contribution to green-house gas emissions (Thomas et al., 2020). Landscape-level emissions were calculated as the sum of emissions across all pixels.

#### 1.3.2. Carbon stocks

Carbon stocks were assessed from a look-up table relating the land cover of each pixel and its estimated contribution potential for carbon storage (Case and Ryan, 2020; Mason et al., 2012; Thomas et al., 2020). Landscape-level carbon stocks were calculated as the sum of carbon stocks across all pixels.

#### 1.3.3. Nitrogen retention

We modelled nitrogen retention using the nutrient delivery ratio model of the InVEST modelling platform (“Natural Capital Project,” 2023) (rinvest package, Stachelek, 2022). This model simulates nutrient movements due to surface flow, representing the steady-state flow of nutrients. The sources of nutrients, and their transport rates across each land use, were estimated for each land cover (Lavorel et al., 2022). Landscape-level service was calculated as the proportion between landscape nitrogen load and nitrogen export across the whole landscape.

#### 1.3.4. Erosion control

We calculated mean annual erosion rates using the Universal Soil Loss Equation (Renard et al., 1991), which is the product of precipitation, slope gradient, slope length, a soil factor and a vegetation factor. The soil and precipitation factors were constant and estimated from assumptions (Lavorel et al., 2022), the slope length and gradient were derived from the topographic model of each landscape, and the vegetation factor was parameterized for each land cover type based on expert assessment. Erosion control was then calculated as minus the mean annual erosion rates.

#### 1.3.5. Crop pollination

The crop pollination model characterizes land covers based on their requirement for pollination and their suitability as habitats for pollinators. We assumed a pollinator dispersal distance of 500 m, i.e. pollinator habitats provided a pollination services to land covers within a 500m radius (Schulp et al., 2014). The landscape-level pollination service indicator was measured as the proportion of cropland supplied by pollination services.

#### 1.3.6. Recreation

We quantified relative landscape attractiveness using a recreation opportunity spectrum approach (Lavorel et al., 2020). The approach combines three components. Areas within 500 m of a watercourse or areas located in elevated were considered attractive. The third component weights each pixel depending on the relative attractiveness of its land cover and the size of patch of that land cover (Lavorel et al., 2022).

#### 1.3.7. Meteorological shelter / Wind protection

Woody elements act as windbreaks, change surrounding temperatures and affect evapotranspiration (Campi et al., 2012), overall providing meteorological shelter in the surrounding landscape. We estimated the level of shelter based on the friction velocity reduction factor F (Vigiak et al., 2003)*, which quantifies the decrease in wind velocity due to a barrier (e.g. hedge or woodland). F depends on the distance from the barrier (in multiples of the height of the barrier) and the barrier porosity, itself depending on the optical porosity, height, and width of the barrier. We assumed heights of 5 m for hedges and shrubs, and 20 m for natural forests, plantations and riparian vegetation; widths of 5 m for hedges, 10 m for riparian vegetation, and 100 m for shrubs, forests and plantations. Finally, we assumed optical porosities of 0.15 for natural forests, 0.2 for plantations, 0.25 for riparian vegetation, and 0.3 for hedges and shrubs (based on the most similar land uses described in* Vigiak et al., 2003).

Wind speed mitigation was measured as the friction velocity reduction factor f_xh_ following Vigiak et al. 2003:

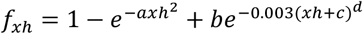

With xh the distance from the barrier, in multiples of barrier heights. A, b, c and d depend on barrier porosity p:

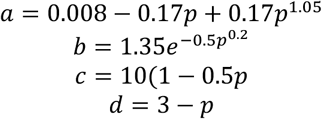

And barrier porosity p is calculated from the barrier optical porosity o, width w, and height h: 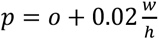 Optical porosity, width and height associated to each type of land use can be found in Table S1 and in the main text.

*F is usually measured* both windward and leeward, and has an asymmetrical distribution around the barrier. However, in the virtual landscapes we assumed wind directionality on the y axis, both directions (north-south, south-north) and the x axis, both directions (east-west, west-east) and thus ignored the windward effect, which is weaker than the leeward one. F was calculated independently for all woody elements and wind directions, and then for each pixel the lowest F (i.e. maximal velocity reduction) was retained. Landscape-level wind protection was measured as minus the sum of F across the non-woody areas of the landscape.

#### 1.3.8. Crop production

We measured crop production as the total amount of energy produced by croplands and grasslands in the landscape. The energy produced by hectare for each land use was obtained by multiplying yield by energy content of each crop (Agreste 2008-2012), weighted by the relative area of the crop in the Isère region in France (Département de l’Isère and Direction Départementale des Territoires, 2019). Crops included winter and spring crops, vegetables and orchards.

### 1.4. Landscape connectivity for different groups

The first three connectivity group were characterized by an affinity for forests and sensitivity to intensive open land uses. The first group had long dispersal abilities and large habitat patch requirement; representative of large carnivores that roam a large distance but are dependent on large patches of high-quality habitat, such as the Eurasian lynx *(Lynx lynx;* hereafter: large carnivores). The second group had a short dispersal range and small patch size requirement, and could use restored riparian forests as well as larger forests. This could represent forest amphibians such as the Yellow-bellied toad (*Bombina variegata,* hereafter: amphibians). The third focal species had a long dispersal range and small patch size requirement (hereafter bats, e.g. Bechstein’s bat, *Myotis bechsteinii*).

We generated the fourth and fifth connectivity groups as specialists with contrasted habitat requirements. The fourth group had a long dispersal range and large patch requirements, but was strictly restricted to forest core areas, avoiding open areas but also forest edges, hedges and linear woody elements. This could represent some forest specific birds, such as the Great spotted woodpecker (*Dendrocopos major*; hereafter: forest specialists). In contrast, the fifth group was an edge specialist, thriving in forest edges and linear woody elements. This can represent tree-nesting bird species that feed on arthropods provided by open habitats such as the European robin (*Erithacus rubecula*; hereafter: edge specialists).

These five groups were associated with positive values either through their intrinsic biodiversity value (all species) or cultural significance (large carnivores, bats, forest specialists). We also considered one connectivity group representing crop weeds such as white goosefoot (*Chenopodium album*), one of the most prominent crop weeds. Because many invasive plant species spread along degraded riverbanks, for this group we not only distinguished restored riparian strips (considered as unfavourable habitat) but also unrestored riverbanks (considered as favourable) whatever the identity of the surrounding land cover.

**Figure S 2.**
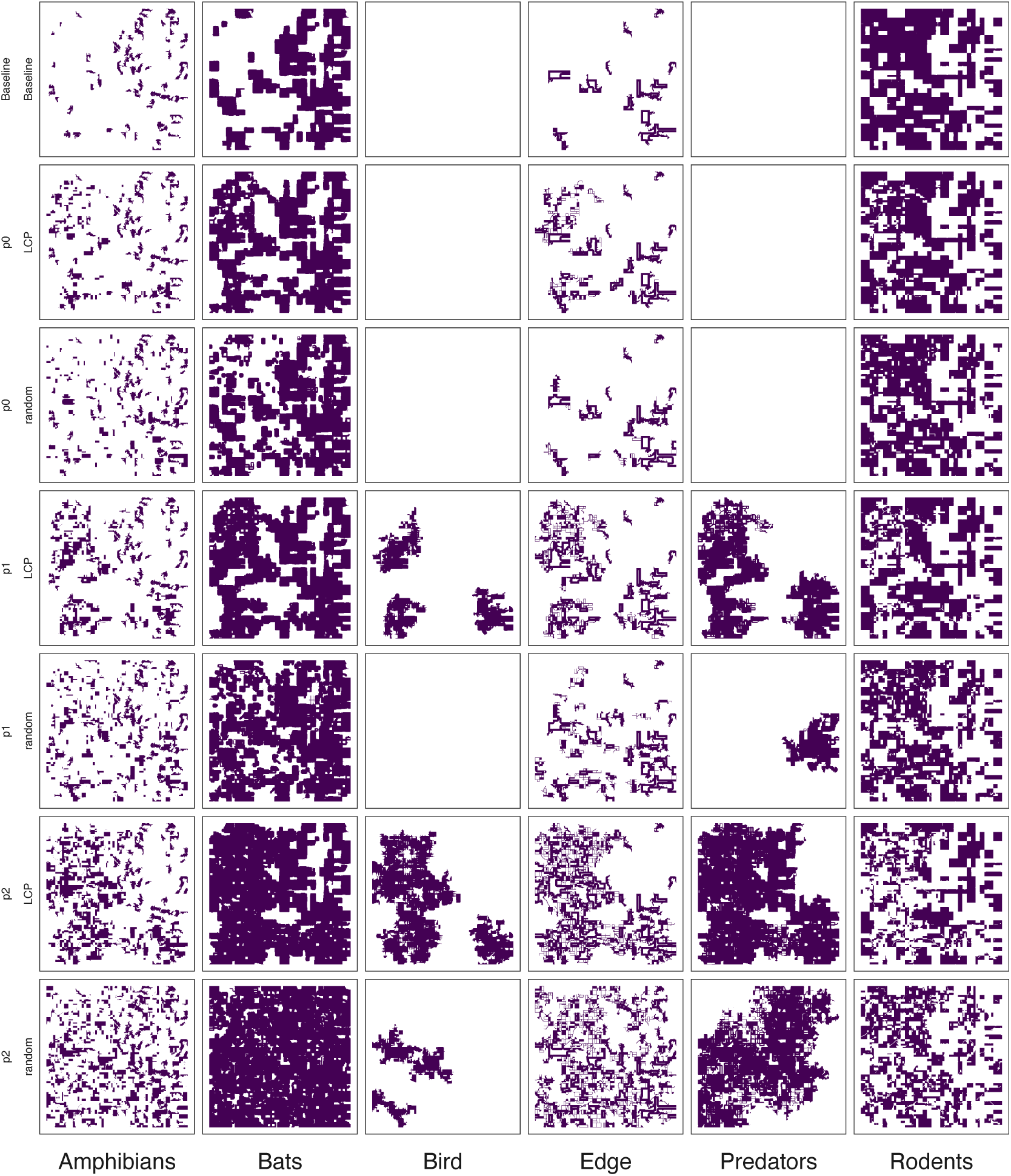
Dispersal kernels for one example landscape for all focal species, proportion restored and two restoration scenarios (random restoration and restoration along corridors). The purple areas show the dispersal kernels, i.e. the area within reach of dispersal from all source patches

## 2. Supplementary results

### 2.1. Variation of landscape metrics across scenarios

**Figure S 3.**
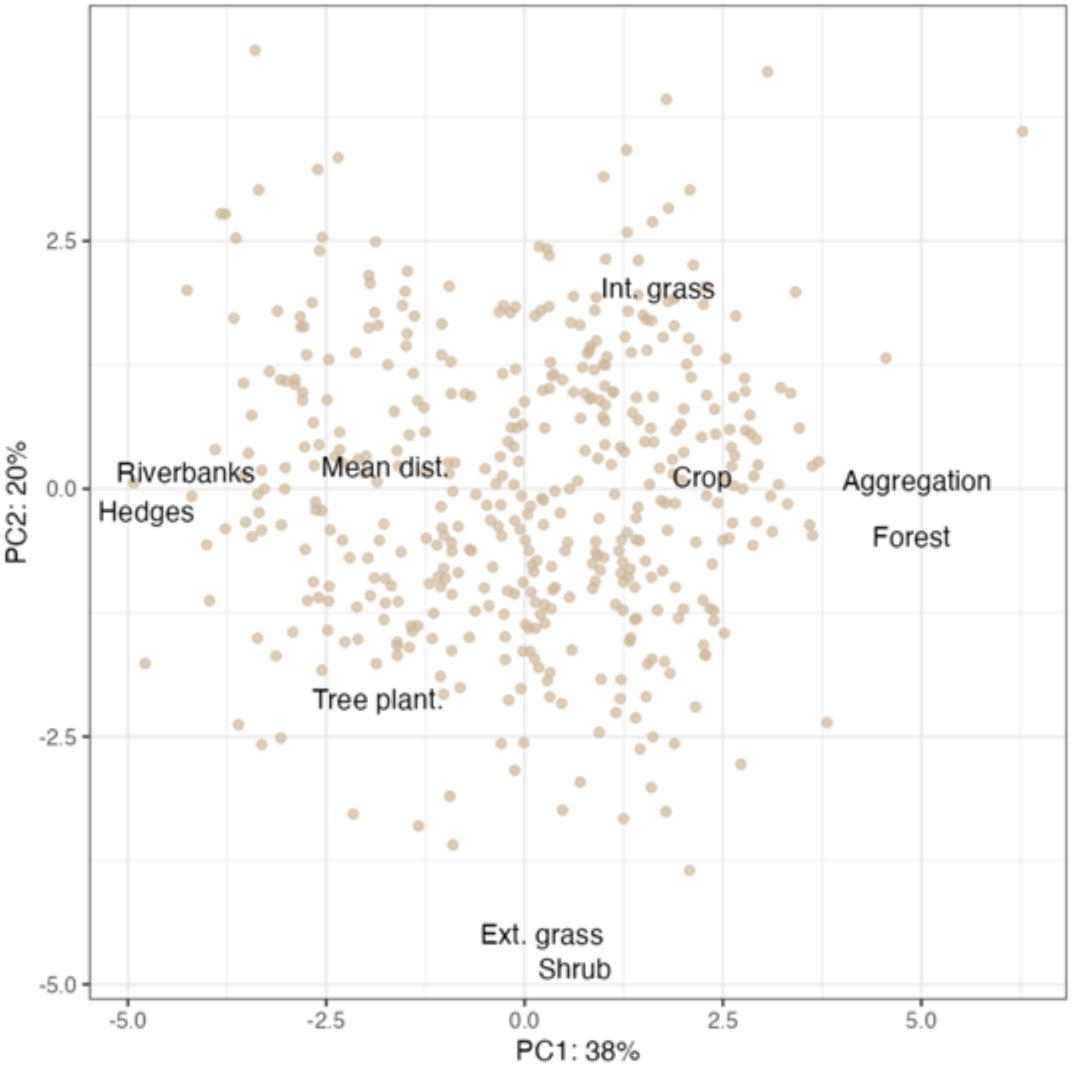
Variation of landscape composition and configuration metrics across landscapes (centred on the proportion restored). Mean dist: mean distance between nearest neighbours of natural cover patches. Aggregation: landscape aggregation index for natural cover. Crop, Tree plantations, Intensive grassland, extensive grassland, Forest, Riverbanks, Hedges: proportion of corresponding land cover in the landscape.

**Figure S 4.**
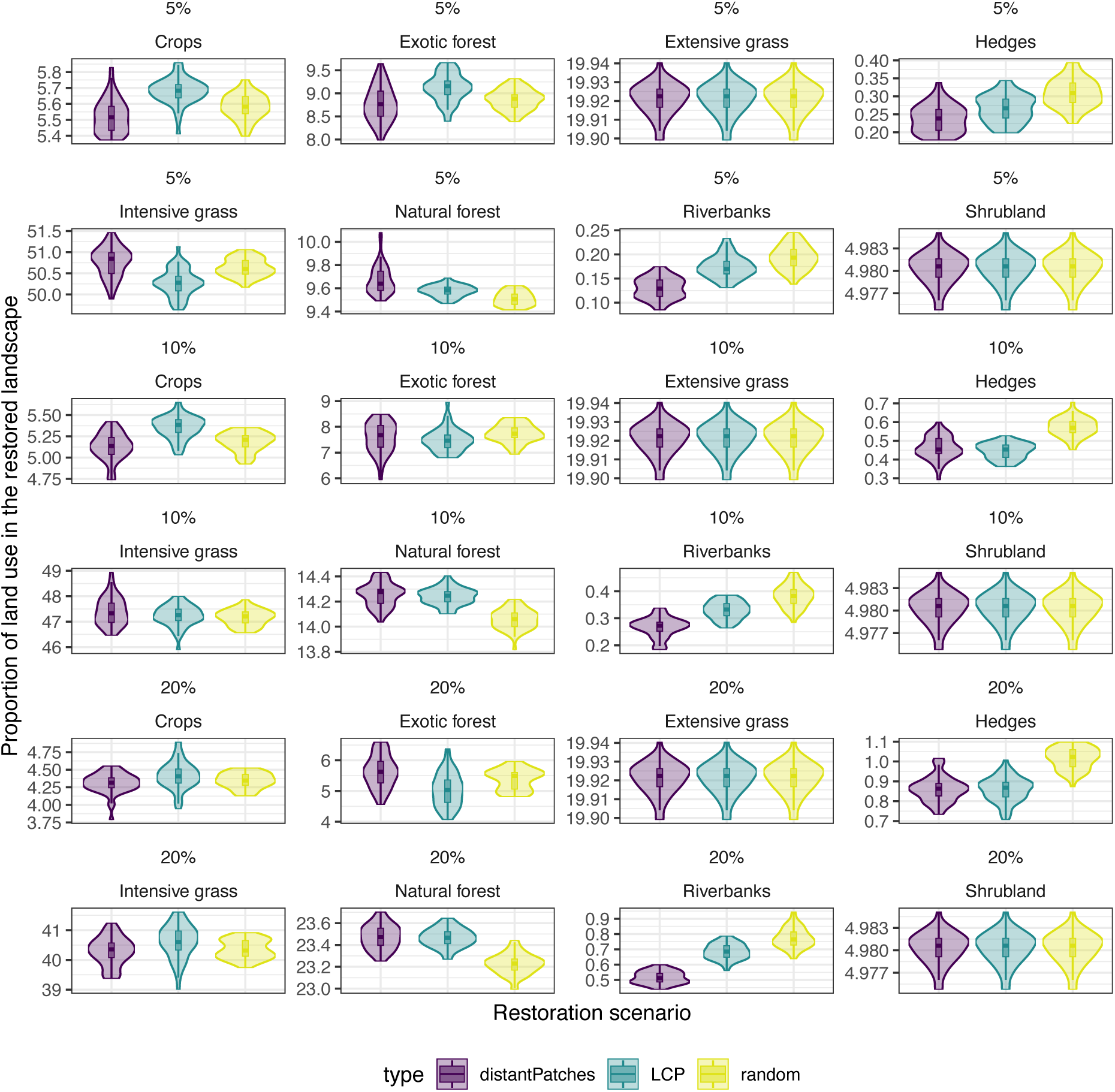
Restoration scenarios outcome vary in term of resulting land use, in particular for the distribution of restored elements.

### 2.2. Variation of individual ecosystem services

**Figure S 5.**
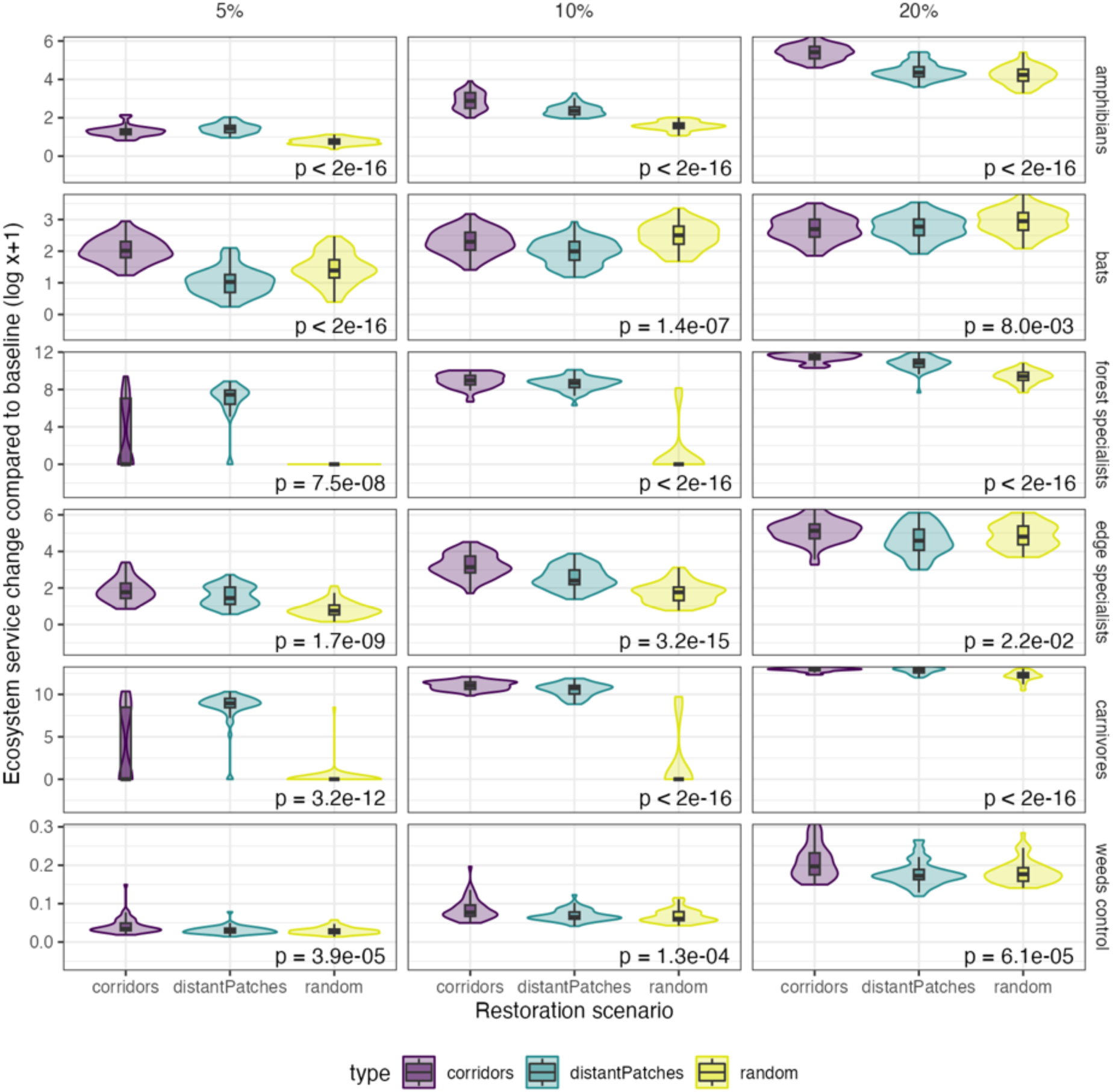
Variation of landscape connectivity for different connectivity groups (shown as log(y +1)) depending on the proportion restored and restoration scenario

**Figure S 6.**
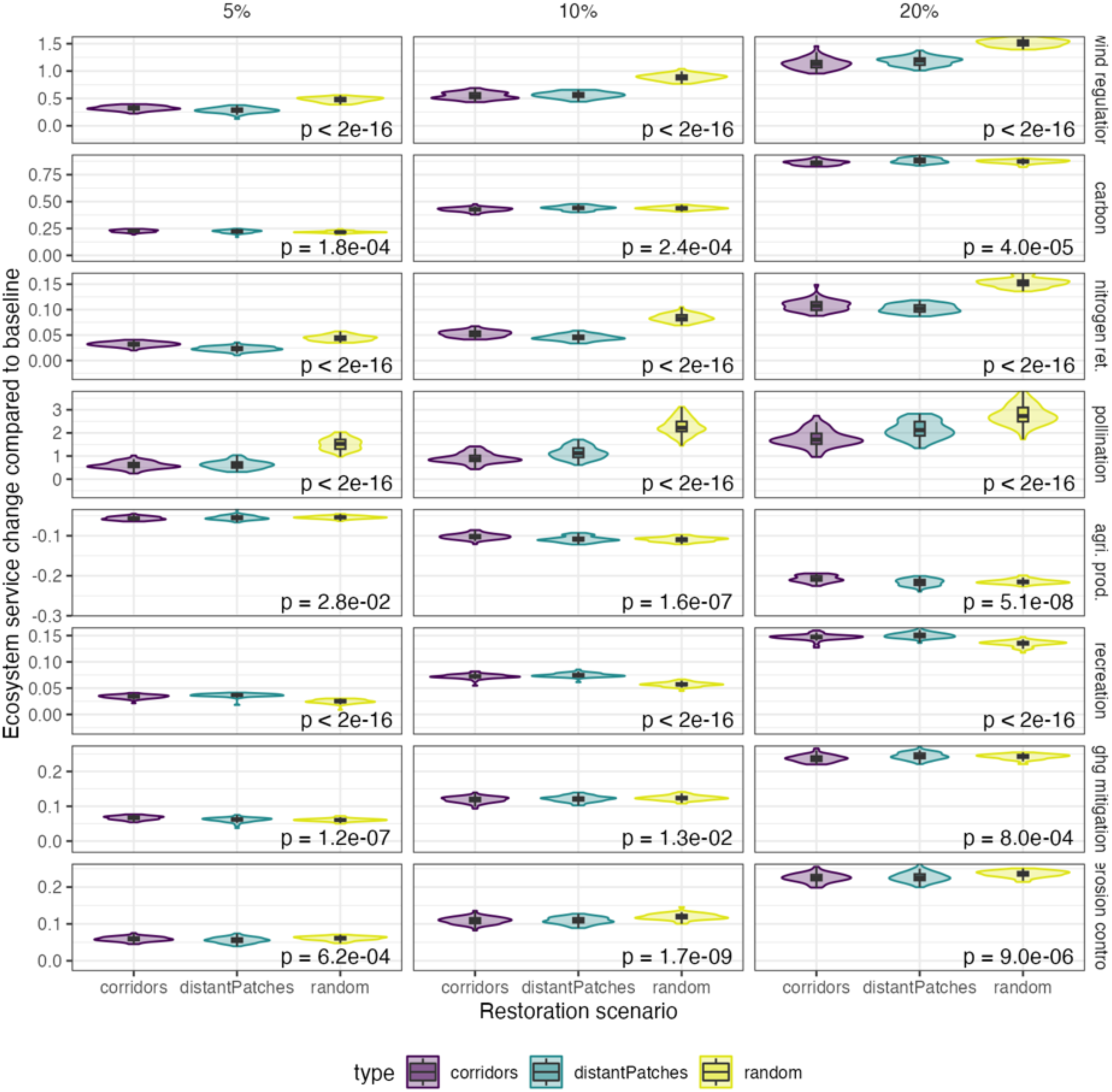
Variation of ecosystem service depending on the proportion restored and restoration scenario

**Figure S 7.**
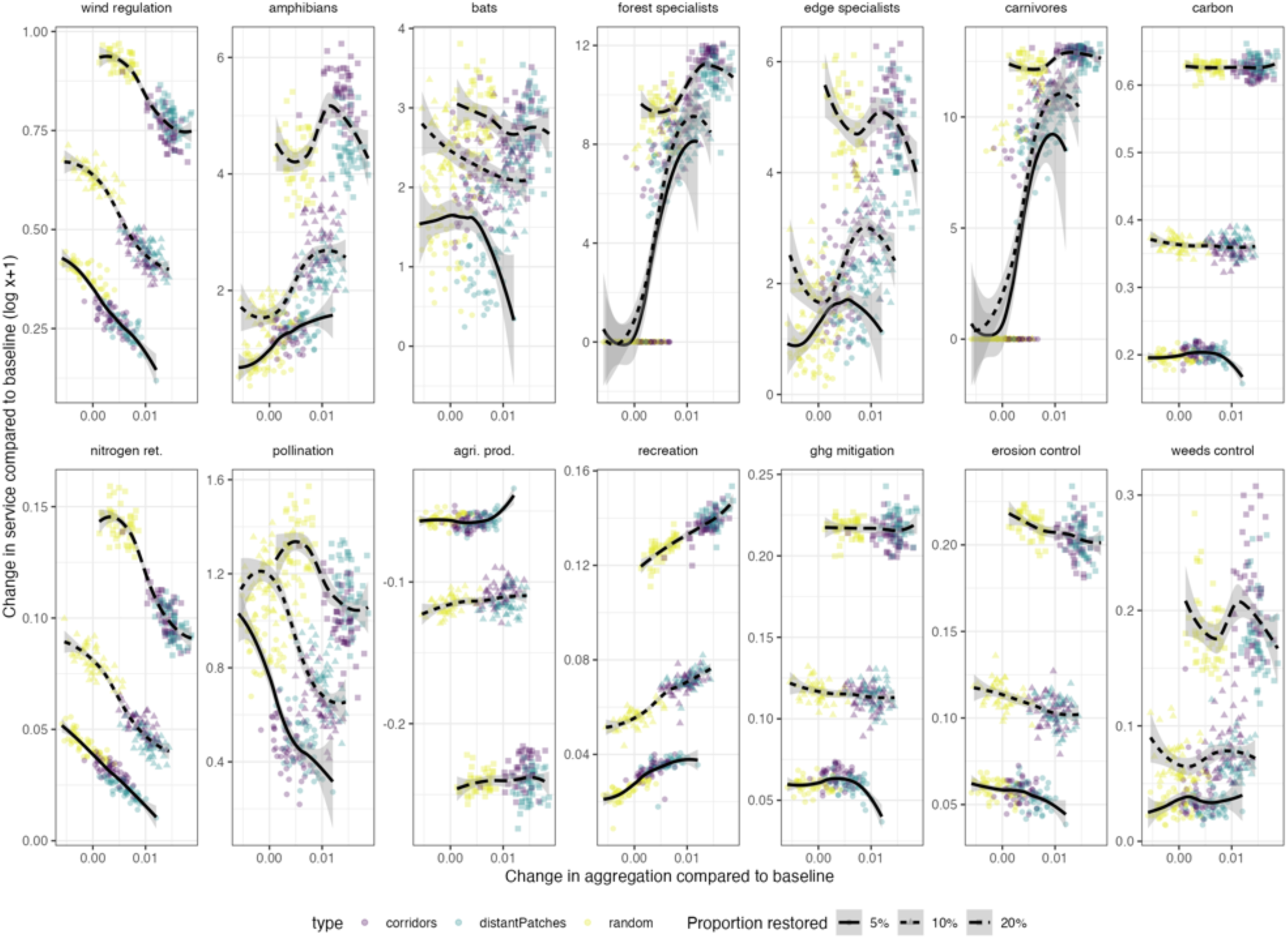
Change in service supply with change in aggregation of natural elements in the landscape compared to the baseline. Shapes and line styles indicate different restoration proportions. Colours indicate different restoration scenarios.

**Table S 2.** Effects of the proportion restored and the aggregation of natural elements in a landscape on each bundle. Showing F and p value for each variable and their interaction, and slope estimates (+ confidence intervals) for the interaction.

| Bundle | Model outputs |  | Proportion restored | Slope per proportion restored (confidence interval) |
| --- | --- | --- | --- | --- |
| Sustainable production | Delta aggregation | F: 737.5 ($p < 10^{-6}$) | 5% | -27.07 (-34.83 - -19.32) |
| | Proportion restored | F: 1738.1 ($p < 10^{-6}$) | 10% | -31.39 (-37.23 - -25.54) |
| | Aggregation*proportion | F: 2.31 ($p = 0.1$) | 20% | -21.84 (-28.3 - -15.37) |
|  | R<sup>2</sup> | 90% |  |  |
| Climate adaptation and mitigation | Delta aggregation | F: 361.3 ($p < 10^{-6}$) | 5% | -25.33 (-63.28 - 12.62) |
| | Proportion restored | F: 486.0 ($p < 10^{-6}$) | 10% | -44.56 (-73.19 - -15.93) |
| | Aggregation*proportion | F: 0.37 ($p = 0.69$) | 20% | -31.24 (-62.89 - 0.41) |
|  | R<sup>2</sup> | 75% |  |  |
| Biodiversity | Delta aggregation | F: 2155.6 ($p < 10^{-6}$) | 5% | 80.86 (55.17 - 106.56) |
| | Proportion restored | F: 944.7 ($p < 10^{-6}$) | 10% | 98.07 (78.69 - 117.45) |
| | Aggregation*proportion | F: 13.2 ($p = 2.7 \cdot 10^{-6}$) | 20% | 24.18 (2.76 - 45.61) |
|  | R<sup>2</sup> | 90% |  |  |
| Cultural value | Delta aggregation | F: 2795.0 ($p < 10^{-6}$) | 5% | 75.48 (52.83 - 98.14) |
| | Proportion restored | F: 1198.9 ($p < 10^{-6}$) | 10% | 85.58 (68.49 - 102.67) |
| | Aggregation*proportion | F: 4.29 ($p = 0.01$) | 20% | 48.19 (29.3 - 67.08) |
|  | R<sup>2</sup> | 92% |  |  |

## 3. Sensitivity analyses: resistance conversion factor

### 3.1. Variation in dispersal kernels

**Figure S 8.**
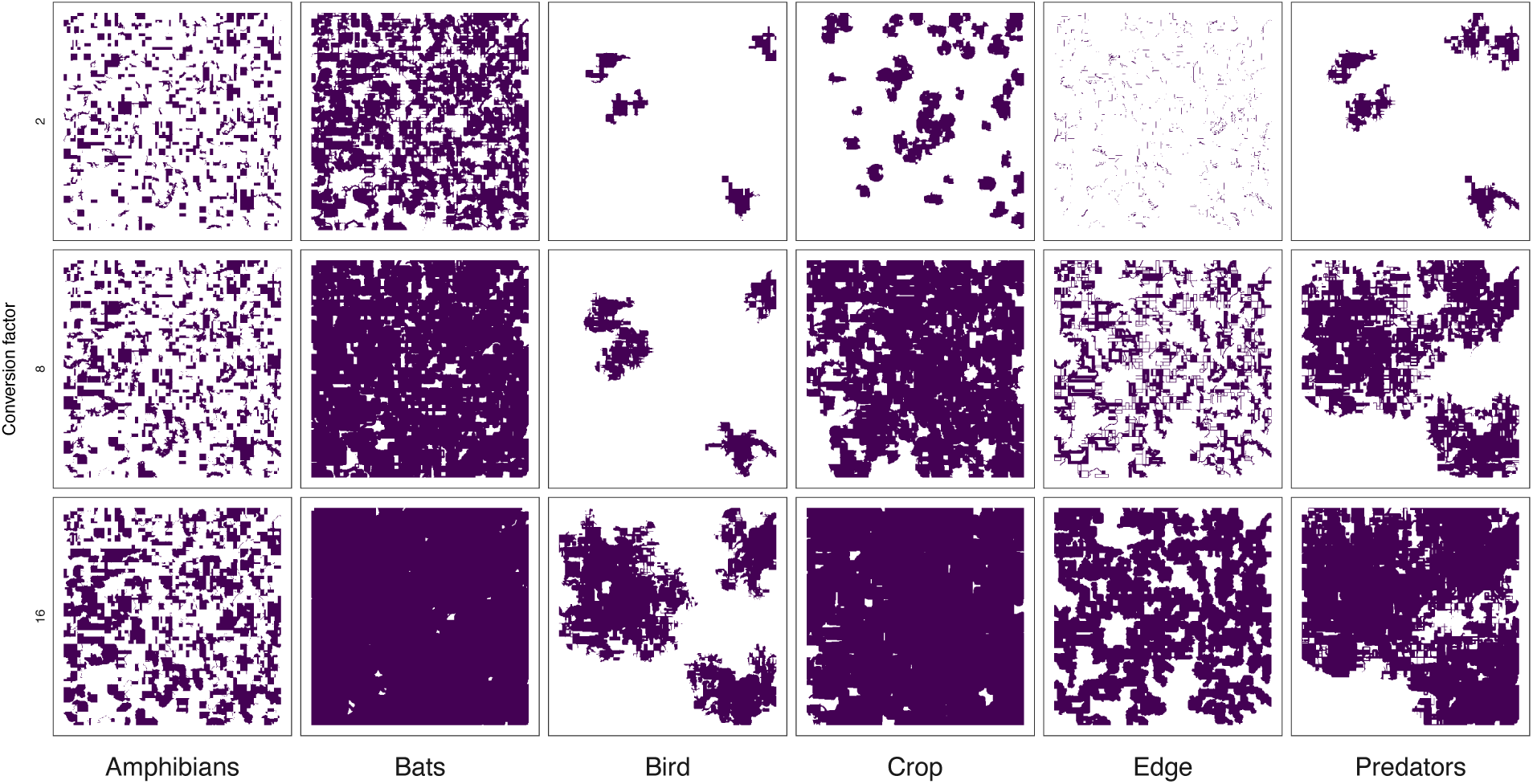
Impact of conversion factor on dispersal kernels for all connectivity groups for an example at 20% restoration (default shown in the main text: 8)

#### 3.1.1. Smaller conversion factor: C = 2

**Figure S 9.**
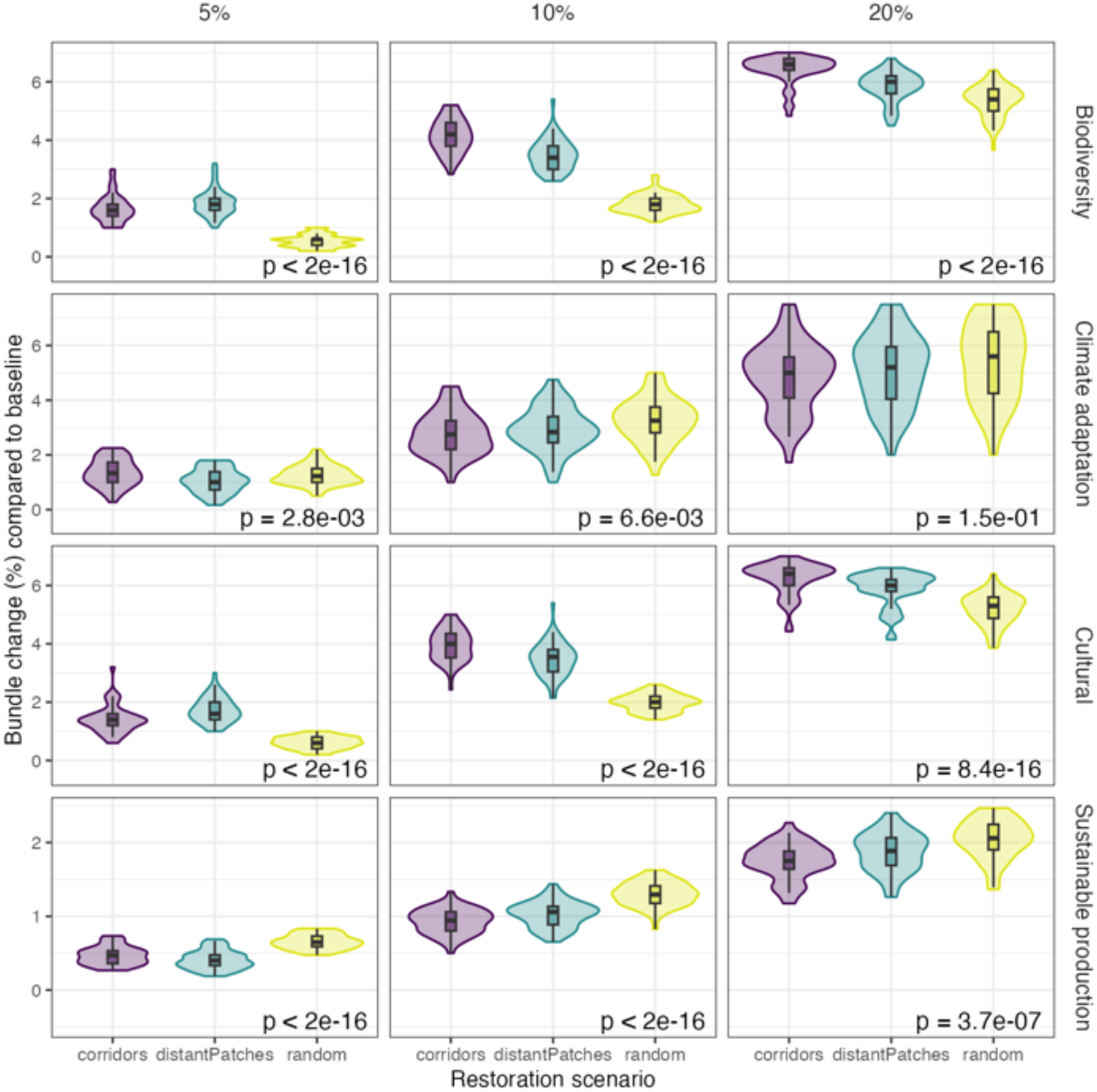
Change in bundle values per restoration scenario and per proportion of area restored. Difference to Fig3: the conversion factor from habitat suitability to resistance was 2 instead of 8 (all other parameters equal)

**Figure S 10.**
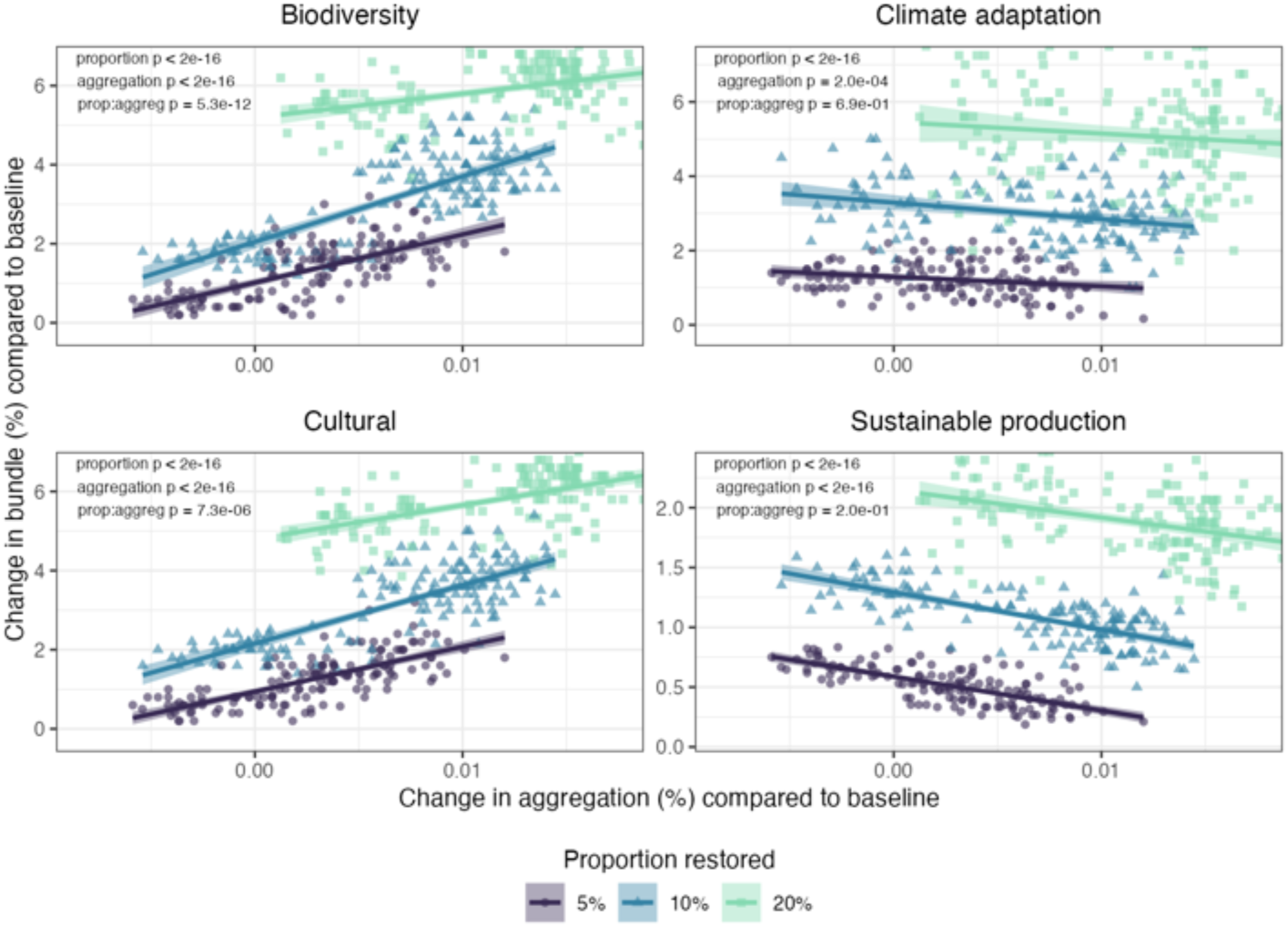
Change in bundle values per change in landscape aggregation. Difference to Fig 4: the conversion factor from habitat suitability to resistance was 2 instead of 8 (all other parameters equal).

#### 3.1.2. Larger conversion factor: C = 16

**Figure S 11.**
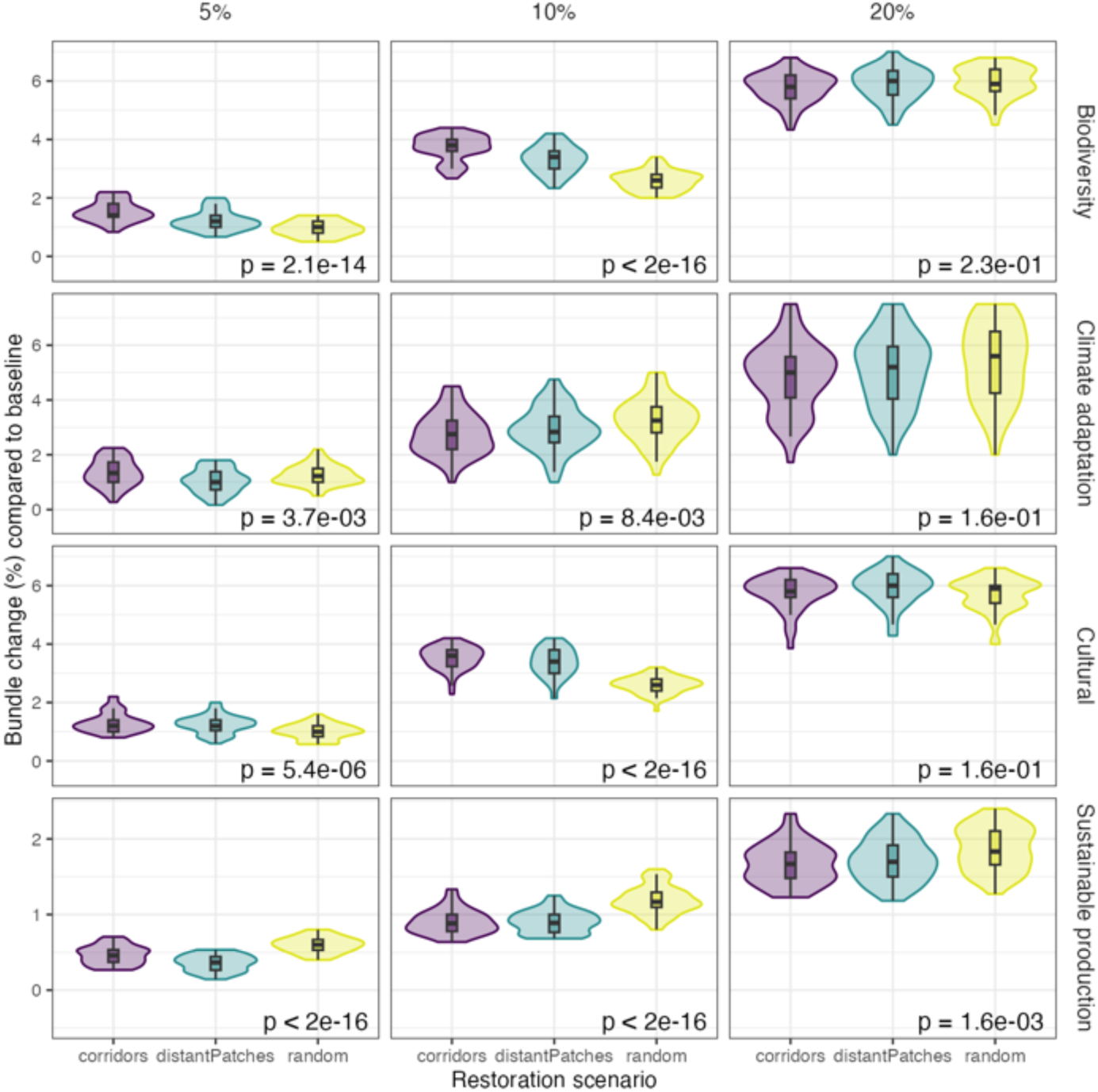
Change in bundle values per restoration scenario and per proportion of area restored. Difference to Fig3: the conversion factor from habitat suitability to resistance was 16 instead of 8 (all other parameters equal)

**Figure S 12.**
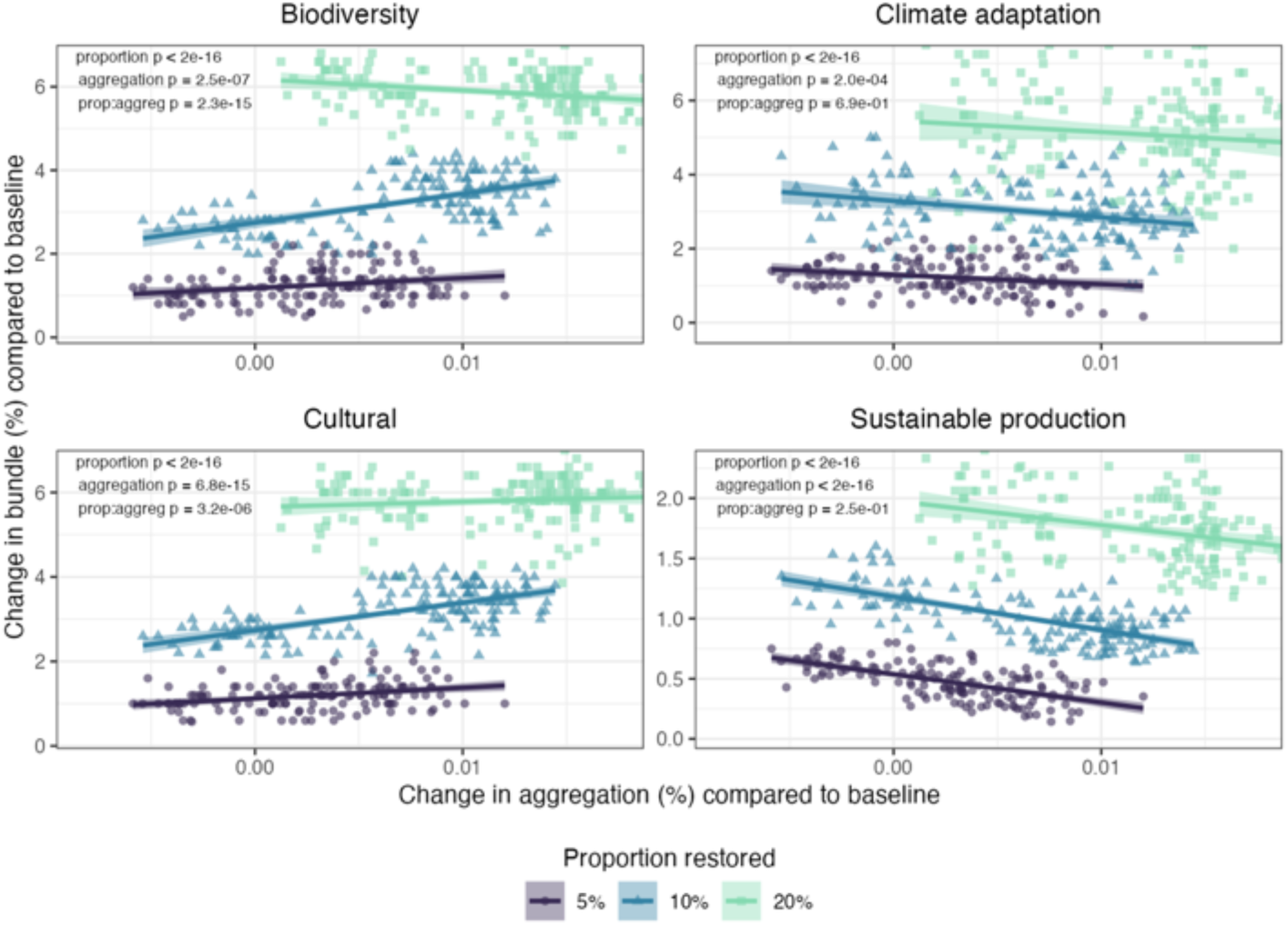
Change in bundle values per change in landscape aggregation. Difference to Fig 4: the conversion factor from habitat suitability to resistance was 16 instead of 8 (all other parameters equal).

### 3.2. Sensitivity analyses: minimal patch sizes in the connectivity analyses

#### 3.2.1. Smaller patch size

**Figure S 13.**
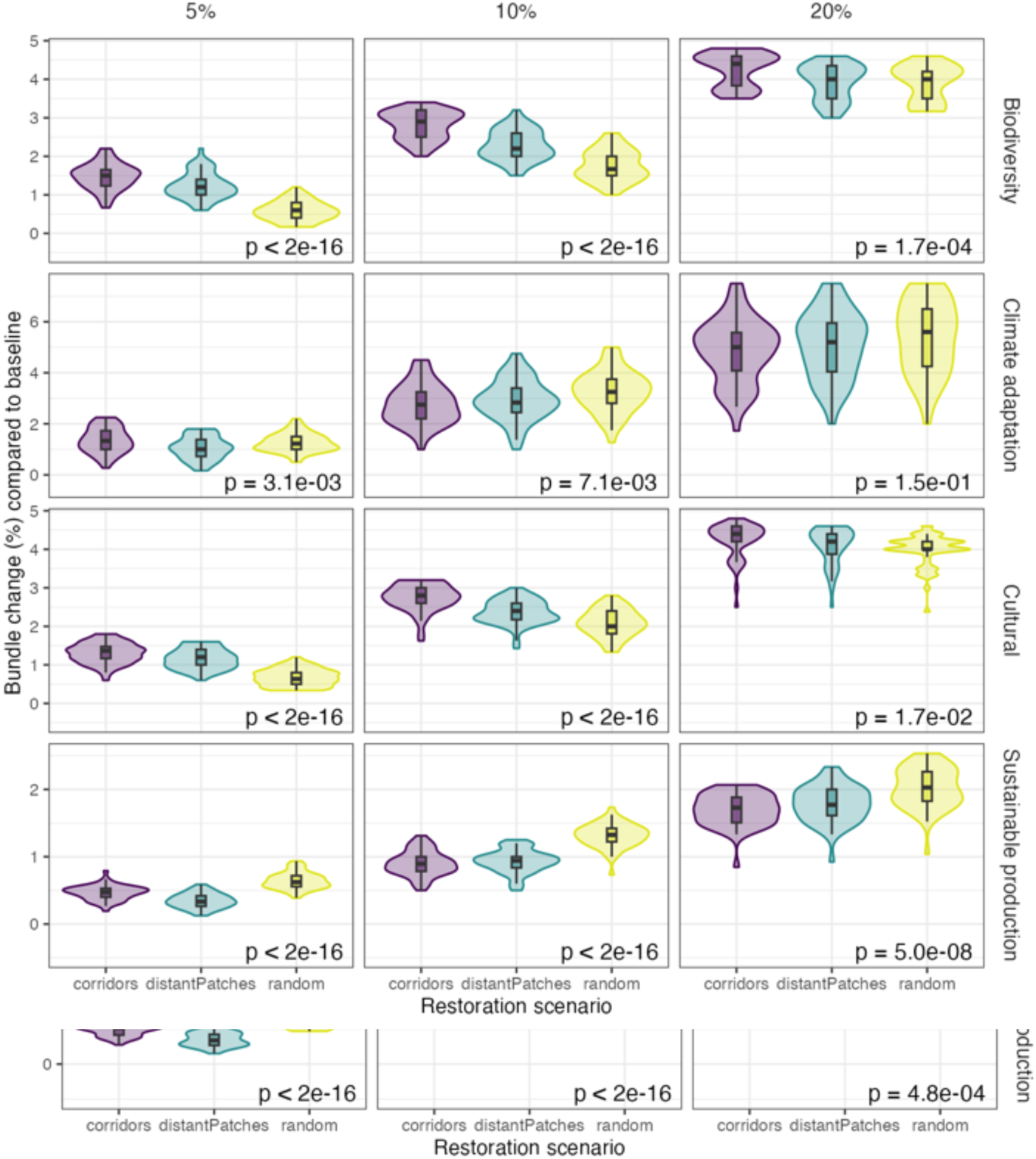
Change in bundle values per restoration scenario and per proportion of area restored. Difference to Fig3: the minimal patch sizes was 0.15 km^2^ and 3km^2^ instead of 0.5 km^2^ and 10 km^2^ for groups with small and large habitat requirements, respectively (all other parameters equal)

**Figure S 14.**
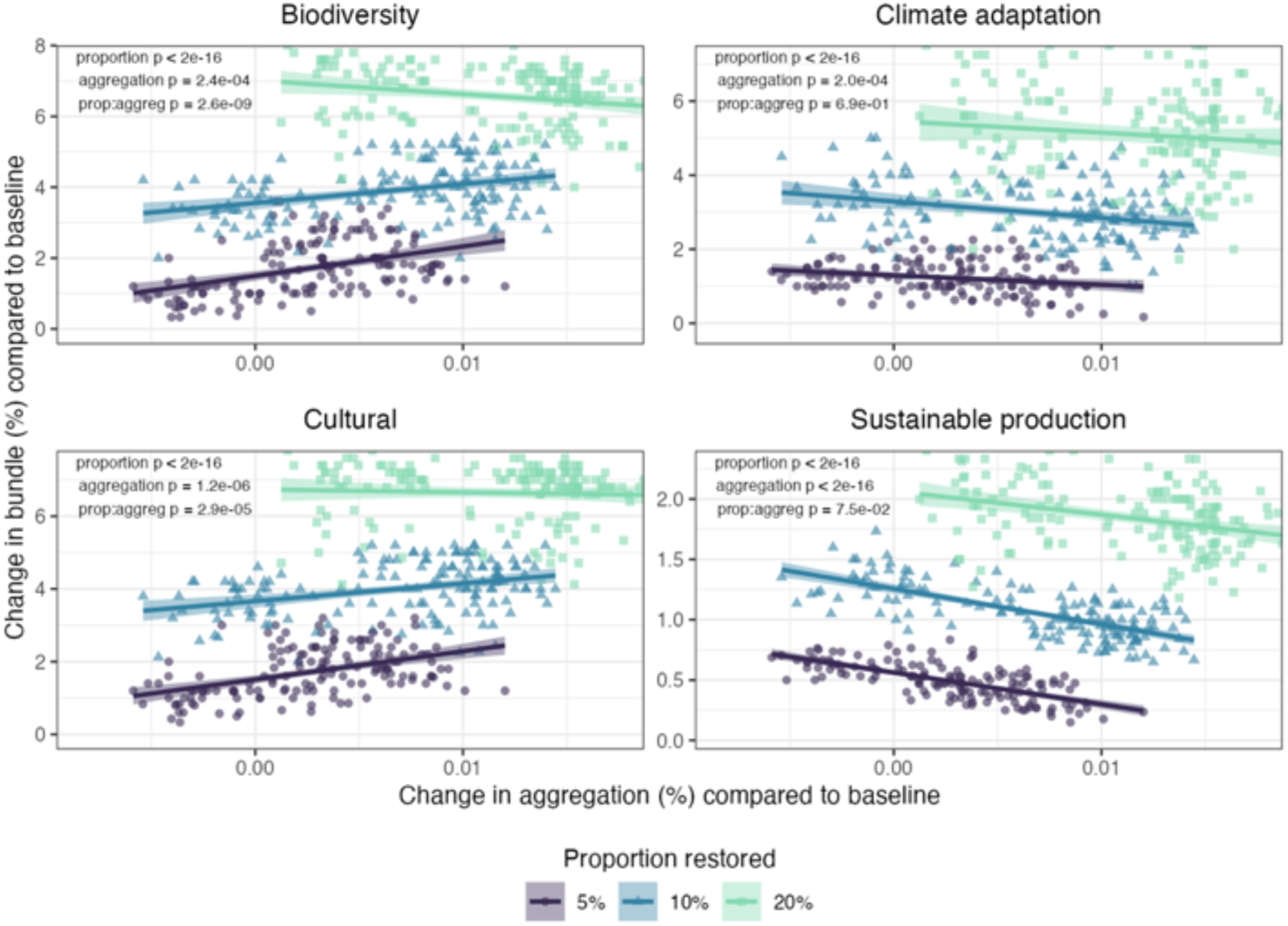
Change in bundle values per change in landscape aggregation. Difference to Fig 4: the minimal patch sizes was 0.15 km^2^ and 3km^2^ instead of 0.5 km^2^ and 10 km^2^ for groups with small and large habitat requirements, respectively (all other parameters equal).

#### 3.2.2. Larger patch size

**Figure S 15.**
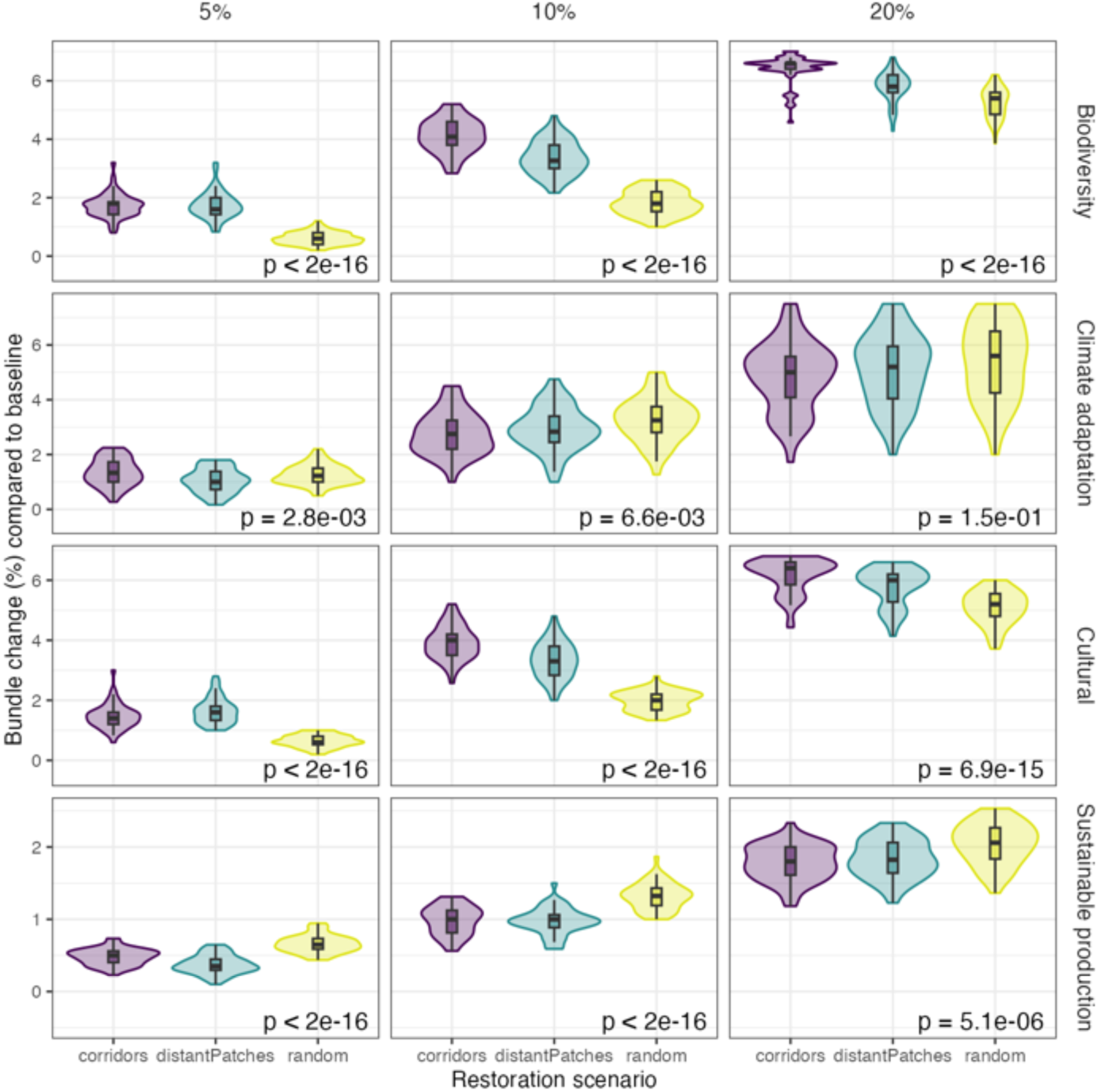
Change in bundle values per restoration scenario and per proportion of area restored. Difference to Fig3: the minimal patch sizes was 1.5 km^2^ and 30km^2^ instead of 0.5 km^2^ and 10 km^2^ for groups with small and large habitat requirements, respectively (all other parameters equal)

**Figure S 16.**
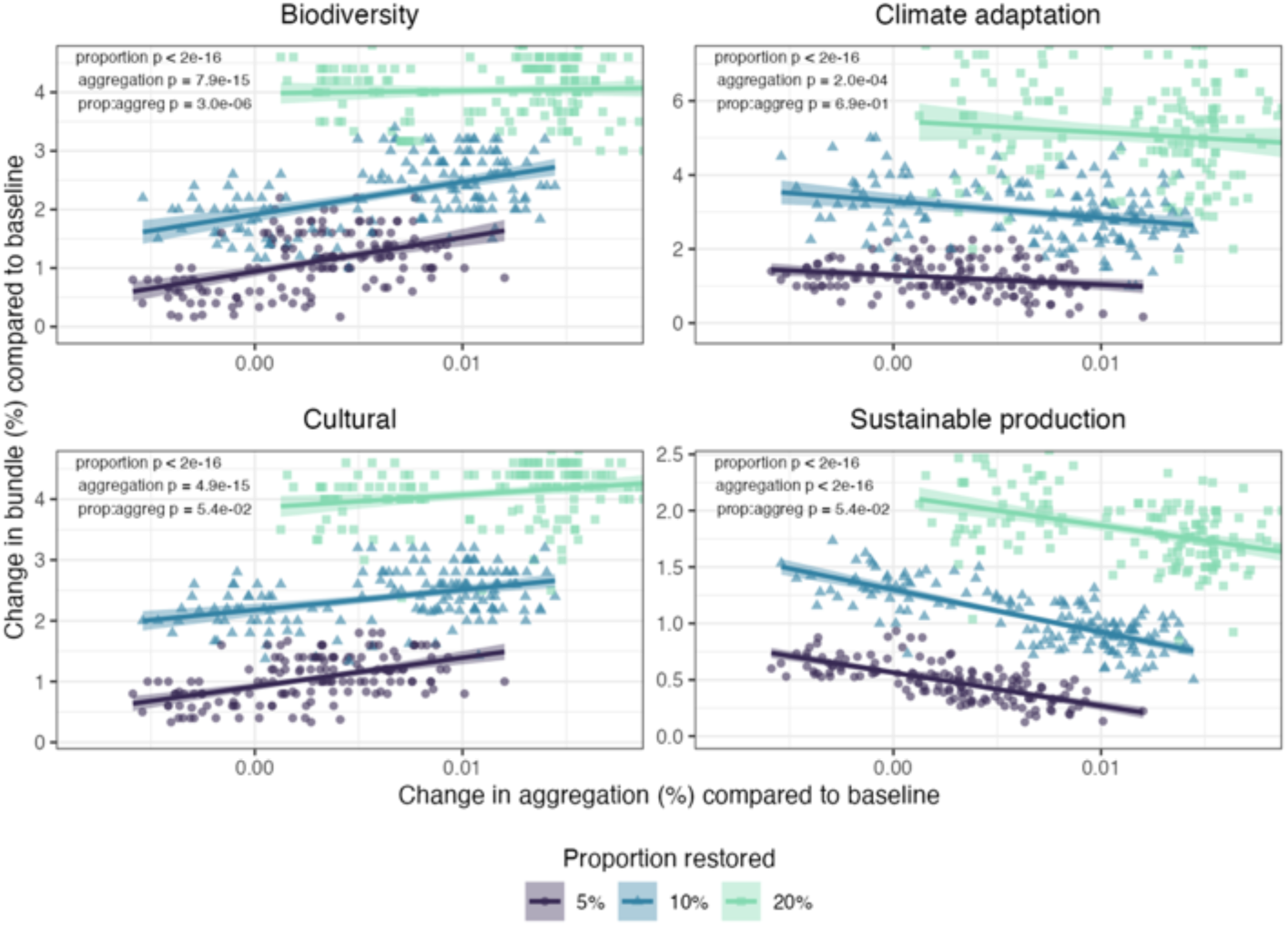
Change in bundle values per change in landscape aggregation. Difference to Fig 4: the minimal patch sizes was 1.5 km^2^ and 30km^2^ instead of 0.5 km^2^ and 10 km^2^ for groups with small and large habitat requirements, respectively (all other parameters equal).

### 3.3. Sensitivity analyses: dispersal distance in the connectivity analyses

#### 3.3.1. Smaller dispersal distance

**Figure S 17.**
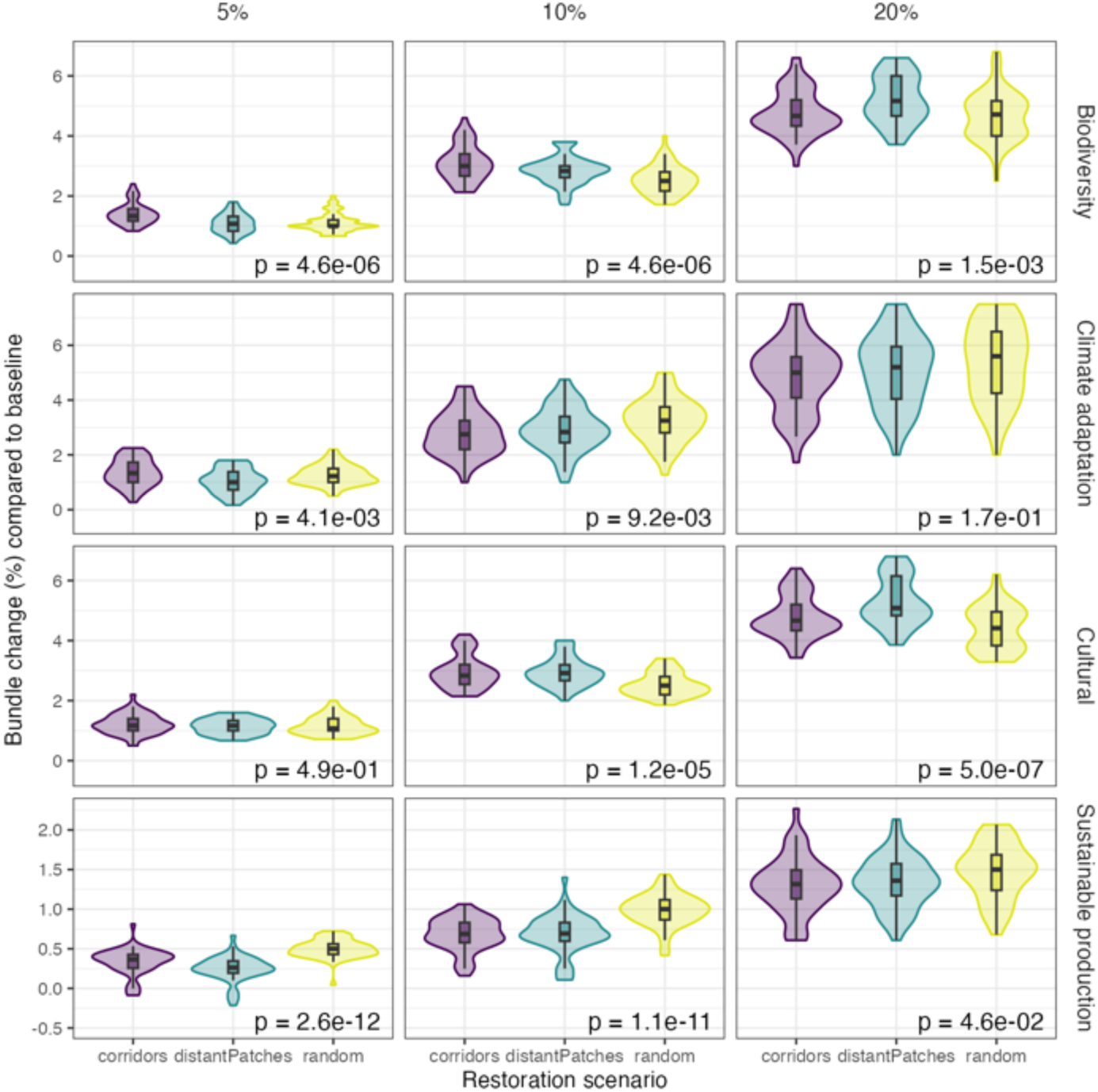
Change in bundle values per restoration scenario and per change in landscape aggregation. Difference to Fig 3: the dispersal distance was 0.25 km and 5km instead of 1 km and 20 km for groups with short and long dispersal, respectively (all other parameters equal)

**Figure S 18.**
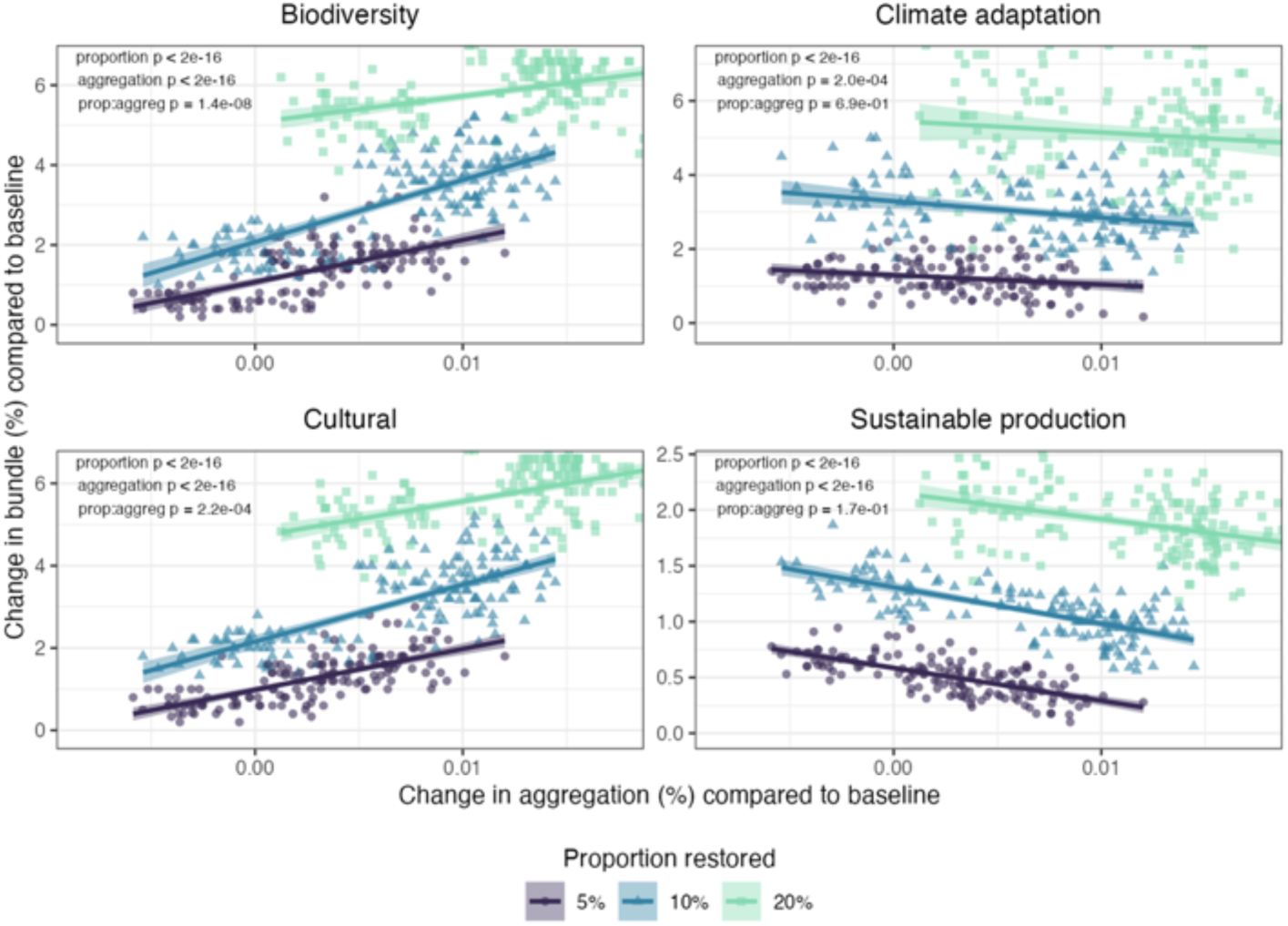
Change in bundle values per change in landscape aggregation. Difference to Fig 4: the dispersal distance was 0.25 km and 5km instead of 1 km and 20 km for groups with short and long dispersal, respectively (all other parameters equal).

#### 3.3.2. Larger dispersal distance

**Figure S 19.**
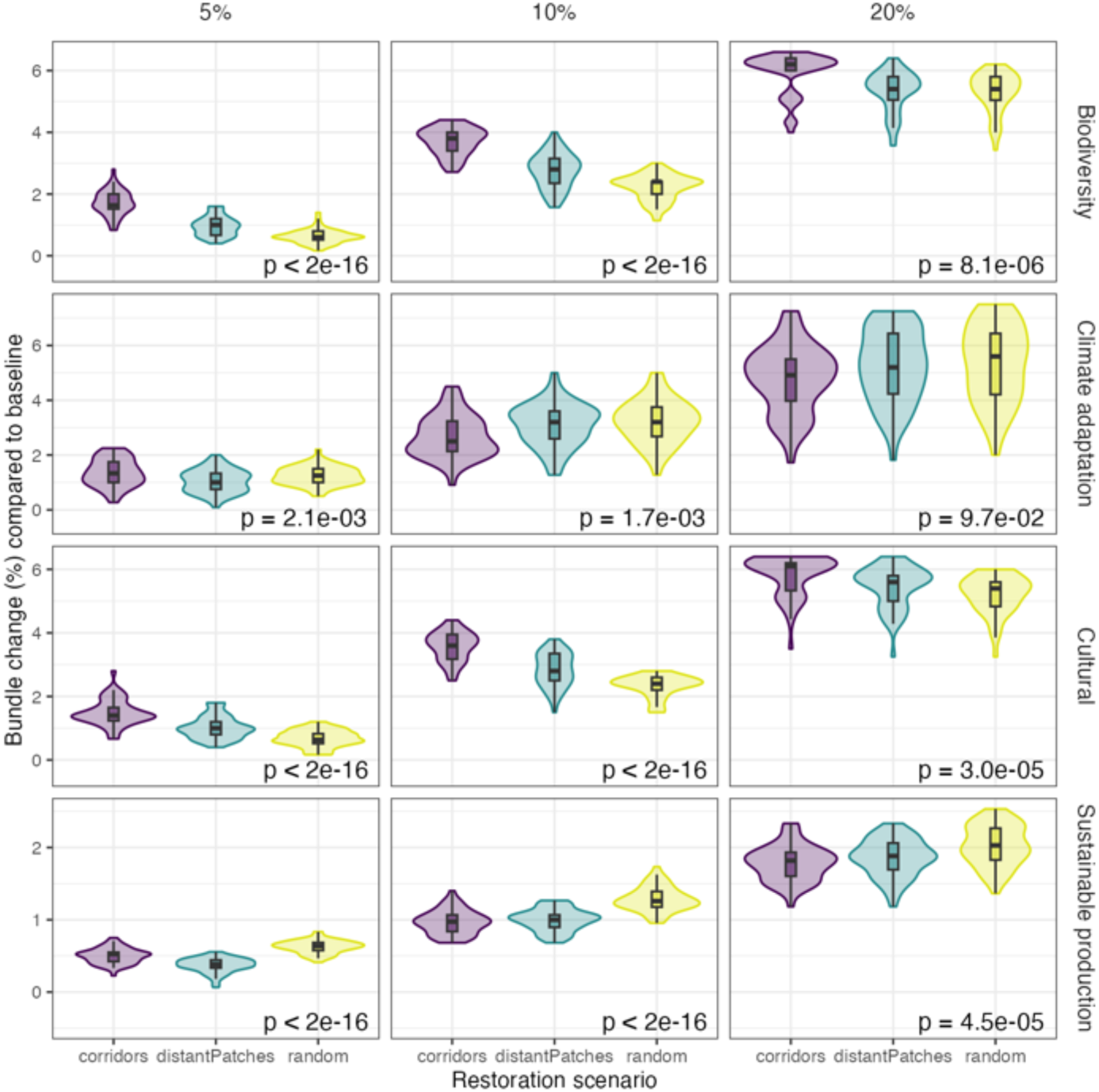
Change in bundle values per restoration scenario and per proportion of area restored. Difference to Fig3: the dispersal distance was 4 km and 80 km instead of 1 km and 20 km for groups with short and long dispersal, respectively (all other parameters equal)

**Figure S 20.**
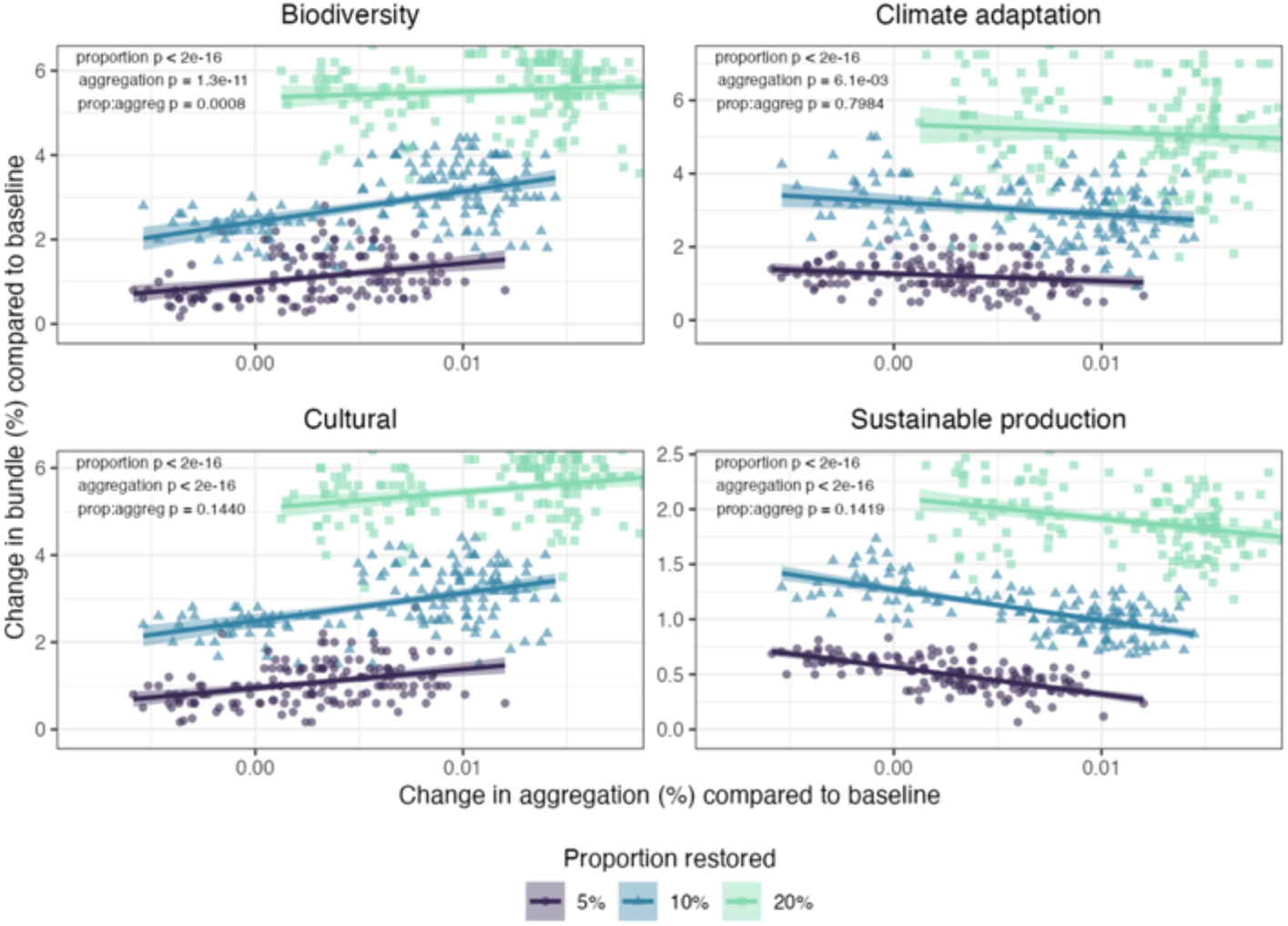
Change in bundle values per change in landscape aggregation. Difference to Fig 4: the dispersal distance was 4 km and 80 km instead of 1 km and 20 km for groups with short and long dispersal, respectively (all other parameters equal).

### 3.4. Sensitivity analyses: target size of restored patches in the distant patches scenarios

#### 3.4.1. Smaller target size

**Figure S 21.**
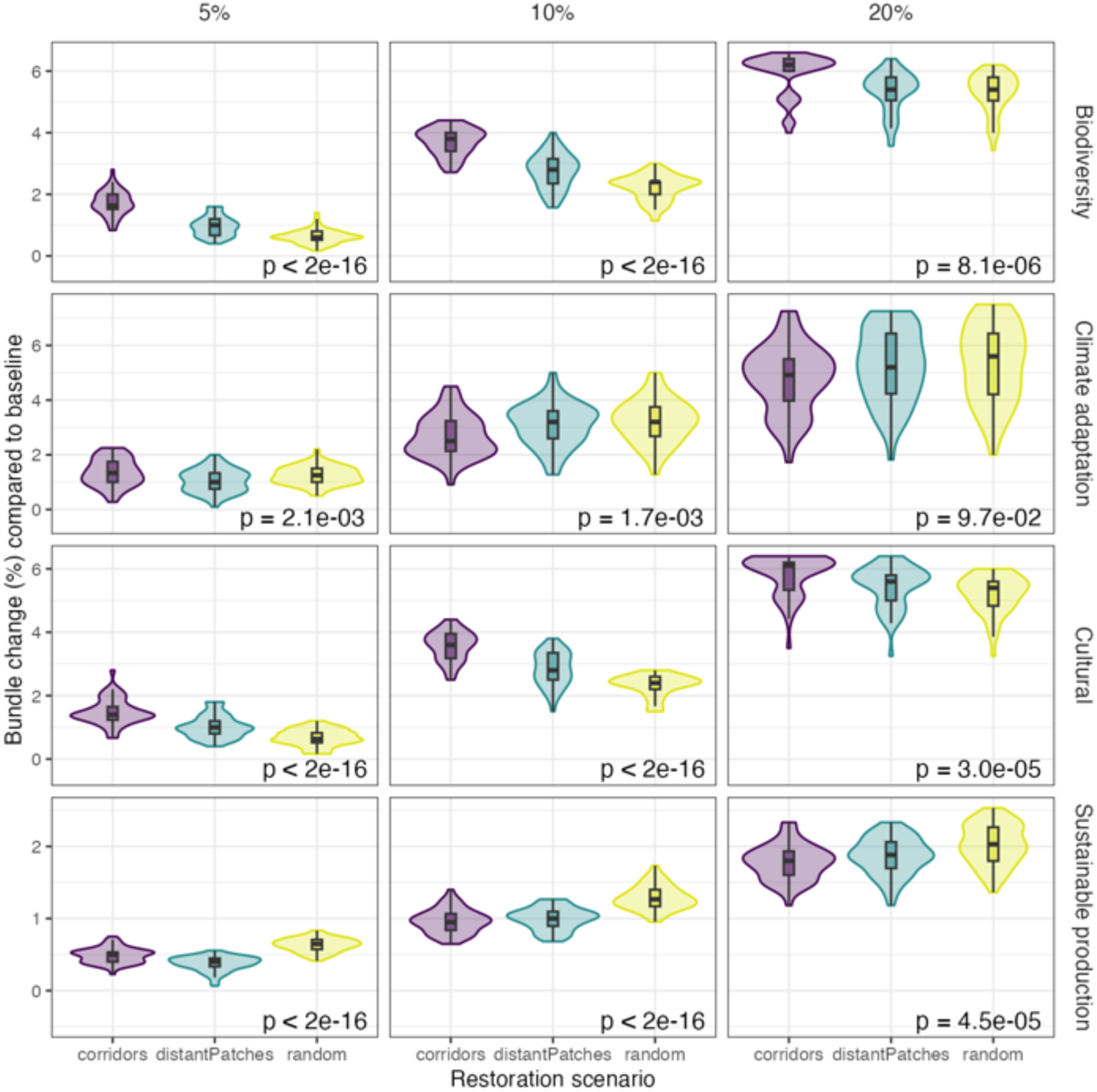
Change in bundle values per restoration scenario and per proportion of area restored. Difference to Fig3: the target size for distant patches was 5 km^2^ instead of 10km^2^ (all other parameters equal)

**Figure S 22.**
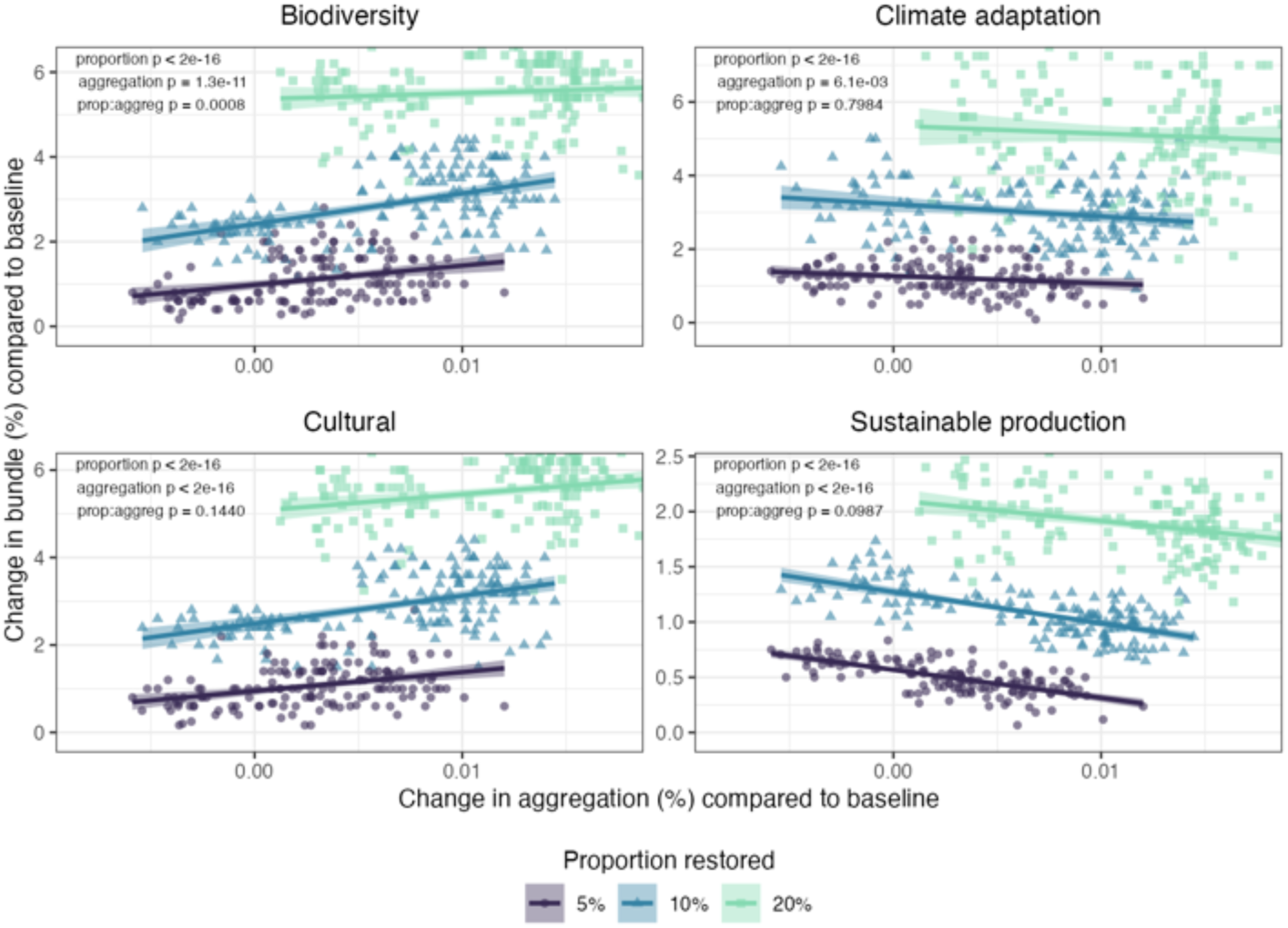
Change in bundle values per change in landscape aggregation. Difference to Fig 4: the target size for distant patches was 5 km^2^ instead of 10km^2^ (all other parameters equal).

#### 3.4.2. Larger target size

**Figure S 23.**
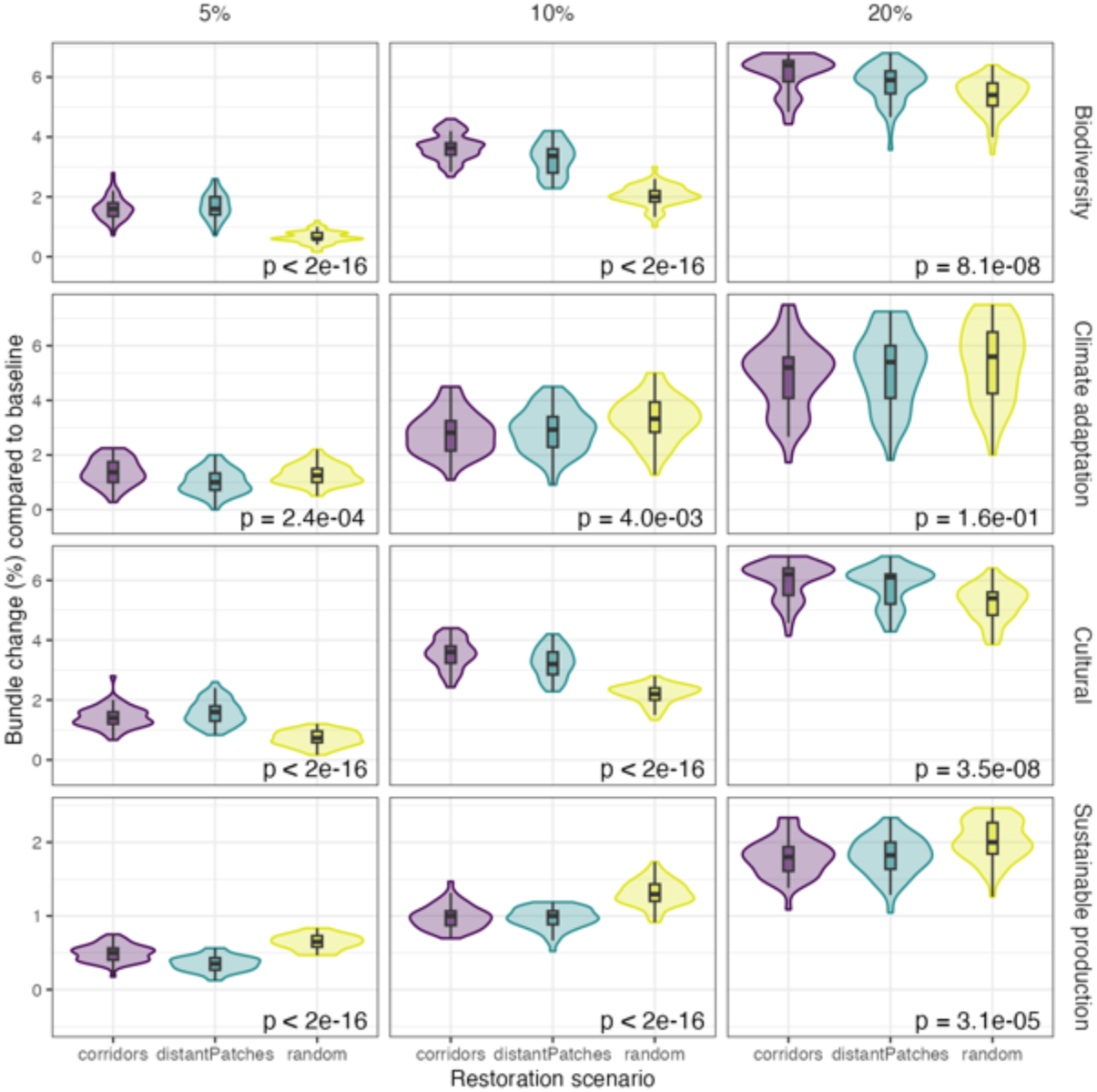
Change in bundle values per restoration scenario and proportion of area restored. Difference to Fig3: the target size for distant patches was 20 km^2^ instead of 10km^2^ (all other parameters equal)

**Figure S 24.**
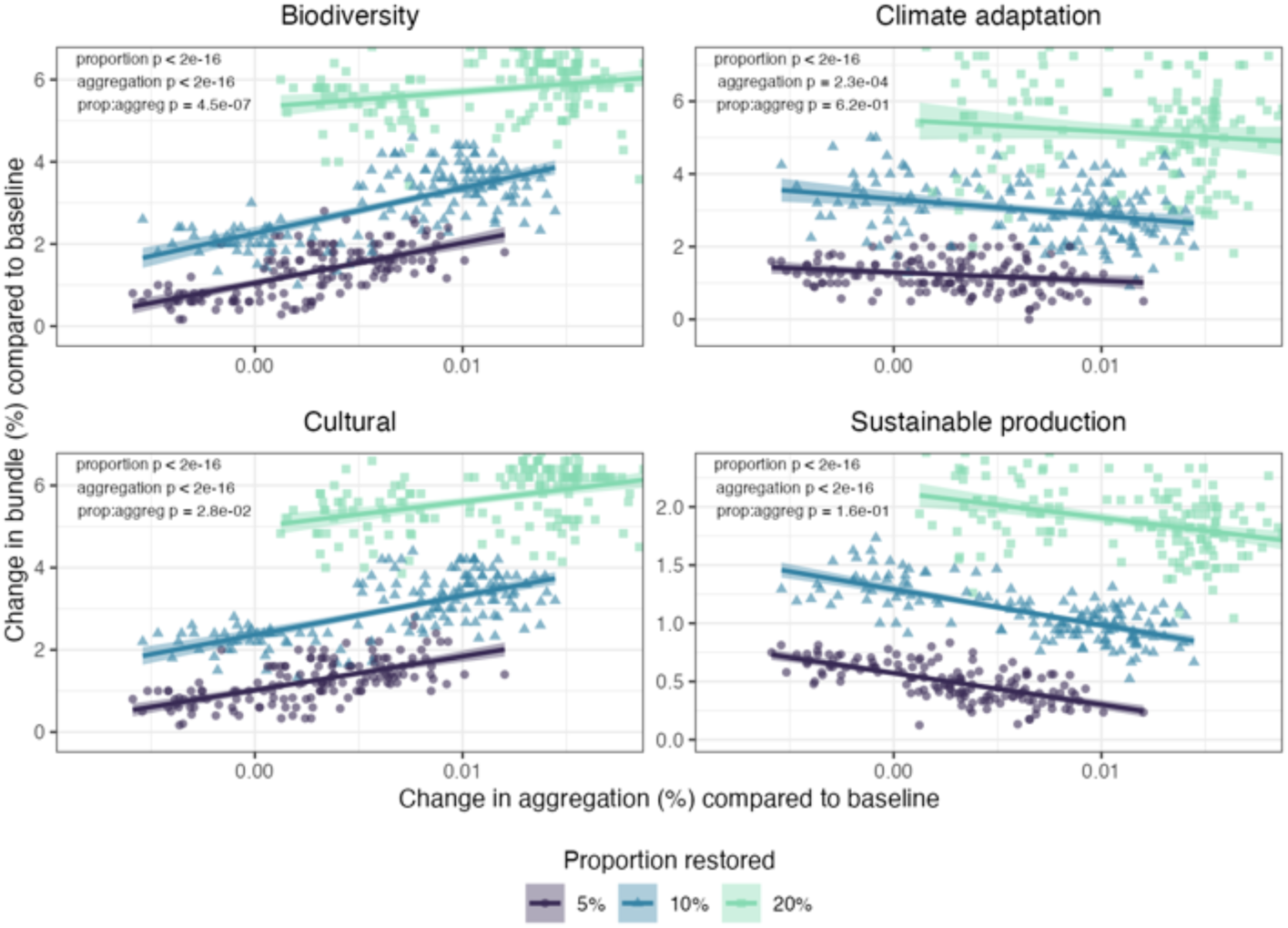
Change in bundle values per change in landscape aggregation. Difference to Fig 4: the target size for distant patches was 20 km^2^ instead of 10km^2^ (all other parameters equal).

## References

Ament, J.M. et al. (2017) ‘Cultural Ecosystem Services in Protected Areas: Understanding Bundles, Trade-Offs, and Synergies’, Conservation Letters, 10(4), pp. 440–450. Available at: 10.1111/conl.12283.

Arroyo-Rodríguez, V. et al. (2020) ‘Designing optimal human-modified landscapes for forest biodiversity conservation’, Ecology Letters, 23(9), pp. 1404–1420. Available at: 10.1111/ele.13535.

Bardgett, R.D. et al. (2021) ‘Combatting global grassland degradation’, Nature Reviews Earth & Environment, 2(10), pp. 720–735. Available at: 10.1038/s43017-021-00207-2.

Barnaud, C. et al. (2023) ‘Participatory research on ecosystem services in the face of disputed values and other uncertainties: A review’, Ecosystem Services, 63, p. 101551. Available at: 10.1016/j.ecoser.2023.101551.

Baró, F., Gómez-Baggethun, E. and Haase, D. (2017) ‘Ecosystem service bundles along the urban-rural gradient: Insights for landscape planning and management’, Ecosystem Services, 24, pp. 147–159. Available at: 10.1016/j.ecoser.2017.02.021.

Boesing, A.L. et al. (2024) ‘Identifying the optimal landscape configuration for landscape multifunctionality’. submitted to Ecosystem Services.

Bommarco, R. et al. (2014) ‘Extinction debt for plants and flower-visiting insects in landscapes with contrasting land use history’, Diversity and Distributions, 20(5), pp. 591–599. Available at: 10.1111/ddi.12187.

Brauman, K.A. et al. (2007) ‘The Nature and Value of Ecosystem Services: An Overview Highlighting Hydrologic Services’, Annual Review of Environment and Resources, 32(1), pp. 67–98. Available at: 10.1146/annurev.energy.32.031306.102758.

Brown, G., Helene Hausner, V. and Lægreid, E. (2015) ‘Physical landscape associations with mapped ecosystem values with implications for spatial value transfer: An empirical study from Norway’, Ecosystem Services, 15, pp. 19–34. Available at: 10.1016/j.ecoser.2015.07.005.

Brumberg, H. et al. (2021) ‘Riparian buffer length is more influential than width on river water quality: A case study in southern Costa Rica’, Journal of Environmental Management, 286, p. 112132. Available at: 10.1016/j.jenvman.2021.112132.

Buchholtz, E.K., Heinrichs, J. and Crist, M. (2023) ‘Landscape and connectivity metrics as a spatial tool to support invasive annual grass management decisions’, Biological Invasions, 25(3), pp. 637–644. Available at: 10.1007/s10530-022-02945-w.

Byczek, C. et al. (2018) ‘Benefits of crowd-sourced GPS information for modelling the recreation ecosystem service’, PLOS ONE, 13(10), p. e0202645. Available at: 10.1371/journal.pone.0202645.

Case, B. and Ryan, C. (2020) An analysis of carbon stocks and net carbon position for New Zealand sheep and beef farmland. Auckland University of Technology. Available at: https://beeflambnz.com/sites/default/files/news-docs/BL_Carbon_report_for_review_final_submit.pdf.

CBD (2022) ‘Decision adopted by the conference of the parties to the Convention on Biological Diversity. Kunming-montreal global biodiversity framework’. Available at: https://www.cbd.int/doc/decisions/cop-15/cop-15-dec-04-en.pdf.

Damschen, E.I. et al. (2019) ‘Ongoing accumulation of plant diversity through habitat connectivity in an 18-year experiment’, Science, 365(6460), pp. 1478–1480. Available at: 10.1126/science.aax8992.

Didham, R.K. et al. (2015) ‘Agricultural Intensification Exacerbates Spillover Effects on Soil Biogeochemistry in Adjacent Forest Remnants’, PLOS ONE, 10(1), p. e0116474. Available at: 10.1371/journal.pone.0116474.

Duarte, G.T. et al. (2018) ‘The effects of landscape patterns on ecosystem services: meta-analyses of landscape services’, Landscape Ecology, 33(8), pp. 1247–1257. Available at: 10.1007/s10980-018-0673-5.

Dudley, N. et al. (2018) ‘The essential role of other effective area-based conservation measures in achieving big bold conservation targets’, Global Ecology and Conservation, 15, p. e00424. Available at: 10.1016/j.gecco.2018.e00424.

Etherington, T.R. (2016) ‘Least-Cost Modelling and Landscape Ecology: Concepts, Applications, and Opportunities’, Current Landscape Ecology Reports, 1(1), pp. 40–53. Available at: 10.1007/s40823-016-0006-9.

Etherington, T.R. et al. (2022) ‘Binary space partitioning generates hierarchical and rectilinear neutral landscape models suitable for human-dominated landscapes’, Landscape Ecology [Preprint]. Available at: 10.1007/s10980-022-01452-6.

Etherington, T.R. (2022) ‘Perlin noise as a hierarchical neutral landscape model’, Web Ecology, 22(1), pp. 1–6. Available at: 10.5194/we-22-1-2022.

Ewers, R.M. and Didham, R.K. (2006) ‘Confounding factors in the detection of species responses to habitat fragmentation’, Biological Reviews, 81(1), pp. 117–142. Available at: 10.1017/S1464793105006949.

Fahrig, L. (2017) ‘Ecological Responses to Habitat Fragmentation Per Se’, Annual Review of Ecology, Evolution, and Systematics, 48(1), pp. 1–23. Available at: 10.1146/annurev-ecolsys-110316-022612.

Farrell, L.E. et al. (2018) ‘Landscape connectivity for bobcat (Lynx rufus) and lynx (Lynx canadensis) in the Northeastern United States’, PLOS ONE, 13(3), p. e0194243. Available at: 10.1371/journal.pone.0194243.

Faure, J., Mouysset, L. and Gaba, S. (2023) ‘Combining incentives with collective action to provide pollination and a bundle of ecosystem services in farmland’, Ecosystem Services, 63, p. 101547. Available at: 10.1016/j.ecoser.2023.101547.

Gardner, R.H. et al. (1992) ‘A Percolation Model of Ecological Flows’, in A.J. Hansen and F. di Castri (eds) Landscape Boundaries: Consequences for Biotic Diversity and Ecological Flows. New York, NY: Springer (Ecological Studies), pp. 259–269. Available at: 10.1007/978-1-4612-2804-2_12.

Garibaldi, L.A. et al. (2023) ‘How to design multifunctional landscapes?’, Journal of Applied Ecology, 60(12), pp. 2521–2527. Available at: 10.1111/1365-2664.14517.

Grass, I. et al. (2019) ‘Land-sharing/-sparing connectivity landscapes for ecosystem services and biodiversity conservation’, People and Nature, 1(2), pp. 262–272. Available at: 10.1002/pan3.21.

Haddad, N.M. et al. (2015) ‘Habitat fragmentation and its lasting impact on Earth’s ecosystems’, Science Advances, 1(2), p. e1500052. Available at: 10.1126/sciadv.1500052.

He, H.S., DeZonia, B.E. and Mladenoff, D.J. (2000) ‘An aggregation index (AI) to quantify spatial patterns of landscapes’, Landscape Ecology, 15(7), pp. 591–601. Available at: 10.1023/A:1008102521322.

IPBES (2019) ‘The global assessment report on Biodiversity and Ecosystem services: summary for policy-makers’, p. 60.

‘IPSI Secretariat’ (2018) The International Partnership for the Satoyama Initiative (IPSI) Information Booklet and 2018 Annual Report. Tokyo: United Nations University Institute for the Advanced Study of Sustainability. Available at: https://collections.unu.edu/eserv/UNU:7425/UNU_IAS_Sayotama_IPSI_2018_AR_v2-min.pdf.

Jaeger, J.A.G. (2000) ‘Landscape division, splitting index, and effective mesh size: new measures of landscape fragmentation’, Landscape Ecology, 15(2), pp. 115–130. Available at: 10.1023/A:1008129329289.

Kärvemo, S. et al. (2017) ‘Forest restoration as a double-edged sword: the conflict between biodiversity conservation and pest control’, Journal of Applied Ecology, 54(6), pp. 1658– 1668. Available at: 10.1111/1365-2664.12905.

Keeley, A.T.H., Beier, P. and Gagnon, J.W. (2016) ‘Estimating landscape resistance from habitat suitability: effects of data source and nonlinearities’, Landscape Ecology, 31(9), pp. 2151–2162. Available at: 10.1007/s10980-016-0387-5.

Lavorel, S. et al. (2019) ‘Mustering the power of ecosystems for adaptation to climate change’, Environmental Science & Policy, 92, pp. 87–97. Available at: 10.1016/j.envsci.2018.11.010.

Lavorel, S. et al. (2020) ‘Interactions between outdoor recreation and iconic terrestrial vertebrates in two French alpine national parks’, Ecosystem Services, 45, p. 101155. Available at: 10.1016/j.ecoser.2020.101155.

Lavorel, S. et al. (2022) ‘Templates for multifunctional landscape design’, Landscape Ecology, 37(3), pp. 913–934. Available at: 10.1007/s10980-021-01377-6.

Le Provost, G. et al. (2022) ‘The supply of multiple ecosystem services requires biodiversity across spatial scales’, Nature Ecology & Evolution, pp. 1–14. Available at: 10.1038/s41559-022-01918-5.

López-Cubillos, S. et al. (2023) ‘Optimal restoration for pollination services increases forest cover while doubling agricultural profits’, PLOS Biology, 21(5), p. e3002107. Available at: 10.1371/journal.pbio.3002107.

Luck, G.W. et al. (2009) ‘Quantifying the Contribution of Organisms to the Provision of Ecosystem Services’, BioScience, 59(3), pp. 223–235. Available at: 10.1525/bio.2009.59.3.7.

Lyons, K.G. et al. (2023) ‘Challenges and opportunities for grassland restoration: A global perspective of best practices in the era of climate change’, Global Ecology and Conservation, 46, p. e02612. Available at: 10.1016/j.gecco.2023.e02612.

Martin, D.A. et al. (2020) ‘Land-use history determines ecosystem services and conservation value in tropical agroforestry’, Conservation Letters, 13(5), p. e12740. Available at: 10.1111/conl.12740.

Mason, N.W.H. et al. (2012) ‘Estimation of current and potential carbon stocks and Kyoto-compliant carbon gain on conservation land’.

McGuire, J.L. et al. (2016) ‘Achieving climate connectivity in a fragmented landscape’, Proceedings of the National Academy of Sciences, 113(26), pp. 7195–7200. Available at: 10.1073/pnas.1602817113.

McRae, B.H. et al. (2008) ‘Using Circuit Theory to Model Connectivity in Ecology, Evolution, and Conservation’, Ecology, 89(10), pp. 2712–2724. Available at: 10.1890/07-1861.1.

Mitchell, M.G.E. et al. (2015) ‘Reframing landscape fragmentation’s effects on ecosystem services’, Trends in Ecology and Evolution, 30(4), pp. 190–198. Available at: 10.1016/j.tree.2015.01.011.

Mitchell, M.G.E., Bennett, E.M. and Gonzalez, A. (2015) ‘Strong and nonlinear effects of fragmentation on ecosystem service provision at multiple scales’, Environmental Research Letters, 10(9), p. 094014. Available at: 10.1088/1748-9326/10/9/094014.

‘Natural Capital Project’ (2023) ‘InVEST’. Stanford University, University of Minnesota, Chinese Academy of Sciences, The Nature Conservancy, World Wildlife Fund, Stockholm Resilience Centre and the Royal Swedish Academy of Sciences.

Neyret, et al. (2023) ‘Landscape management strategies for multifunctionality and social equity’, Nature Sustainability [Preprint]. Available at: 10.1038/s41893-022-01045-w.

Nicholson, E. et al. (2021) ‘Scientific foundations for an ecosystem goal, milestones and indicators for the post-2020 global biodiversity framework’, Nature Ecology & Evolution, 5(10), pp. 1338–1349. Available at: 10.1038/s41559-021-01538-5.

Oehri, J. et al. (2020) ‘Terrestrial land-cover type richness is positively linked to landscape-level functioning’, Nature Communications, 11(1), p. 154. Available at: 10.1038/s41467-019-14002-7.

Ortega, U. et al. (2023) ‘Identifying a green infrastructure to prioritise areas for restoration to enhance the landscape connectivity and the provision of ecosystem services’, Landscape Ecology [Preprint]. Available at: 10.1007/s10980-023-01789-6.

Peter, S. et al. (2022) ‘Cultural Worldviews Consistently Explain Bundles of Ecosystem Serivce Prioritisation Across Rural Germany’, People and Nature, 4(1), pp. 218–230. Available at: DOI: 10.1002/pan3.10277.

van der Plas, F. et al. (2019) ‘Towards the development of general rules describing landscape heterogeneity-multifunctionality relationships’, Journal of Applied Ecology. Edited by E. Nichols, 56(1), pp. 168–179. Available at: 10.1111/1365-2664.13260.

Polyakov, M. et al. (2023) ‘Joining the dots versus growing the blobs: Evaluating spatial targeting strategies for ecological restoration’, Ecological Economics, 204, p. 107671. Available at: 10.1016/j.ecolecon.2022.107671.

Qiu, J. (2019) ‘Effects of Landscape Pattern on Pollination, Pest Control, Water Quality, Flood Regulation, and Cultural Ecosystem Services: a Literature Review and Future Research Prospects’, Current Landscape Ecology Reports, 4(4), pp. 113–124. Available at: 10.1007/s40823-019-00045-5.

Renard, K.G. et al. (1991) ‘RUSLE: Revised universal soil loss equation’, Journal of Soil and Water Conservation [Preprint].

Rey Benayas, J.M. and Bullock, J.M. (2012) ‘Restoration of Biodiversity and Ecosystem Services on Agricultural Land’, Ecosystems, 15(6), pp. 883–899. Available at: 10.1007/s10021-012-9552-0.

Richards, D. et al. (2024) ‘The Importance of Spatial Configuration When Restoring Intensive Production Landscapes for Biodiversity and Ecosystem Service Multifunctionality’, Land, 13(4), p. 460. Available at: 10.3390/land13040460.

Richards, D.R. et al. (2018) ‘Impacts of habitat heterogeneity on the provision of multiple ecosystem services in a temperate floodplain’, Basic and Applied Ecology, 29, pp. 32–43. Available at: 10.1016/j.baae.2018.02.012.

Sandberg, M.S., Regina Lindborg, Simon Jakobsson, Mattias (2016) ‘How to bring historical forms into the future?: An exploration of Swedish semi-natural grasslands’, in Nature, Temporality and Environmental Management. Routledge.

Saura, S. et al. (2017) ‘Protected areas in the world’s ecoregions: How well connected are they?’, Ecological Indicators, 76, pp. 144–158. Available at: 10.1016/j.ecolind.2016.12.047.

Schulp, C.J.E., Lautenbach, S. and Verburg, P.H. (2014) ‘Quantifying and mapping ecosystem services: Demand and supply of pollination in the European Union’, Ecological Indicators, 36, pp. 131–141. Available at: 10.1016/j.ecolind.2013.07.014.

Simpkins, C.E. et al. (2018) ‘Assessing the performance of common landscape connectivity metrics using a virtual ecologist approach’, Ecological Modelling, 367, pp. 13–23. Available at: 10.1016/j.ecolmodel.2017.11.001.

Sirami, C. et al. (2019) ‘Increasing crop heterogeneity enhances multitrophic diversity across agricultural regions’, Proceedings of the National Academy of Sciences, 116(33), pp. 16442– 16447. Available at: 10.1073/pnas.1906419116.

Smith, F.P. et al. (2013) ‘Maximizing retention of native biodiversity in Australian agricultural landscapes—The 10:20:40:30 guidelines’, Agriculture, Ecosystems & Environment, 166, pp. 35–45. Available at: 10.1016/j.agee.2012.01.014.

Thomas, S. et al. (2021) Evaluation of profitability and future potential for low emission productive uses of land that is currently used for livestock. New Zealand Minsitry of Primary Industries. Available at: https://www.mpi.govt.nz/dmsdocument/45847-Evaluation-of-profitability-and-future-potential-for-low-emission-productive-uses-of-land-that-is-currently-used-for-livestock-Technical-report.

Thomas, Z. and Abbott, B.W. (2018) ‘Hedgerows reduce nitrate flux at hillslope and catchment scales via root uptake and secondary effects’, Journal of Contaminant Hydrology, 215, pp. 51–61. Available at: 10.1016/j.jconhyd.2018.07.002.

Tischendorf, L. and Fahrig, L. (2000) ‘On the usage and measurement of landscape connectivity’, Oikos, 90(1), pp. 7–19. Available at: 10.1034/j.1600-0706.2000.900102.x.

Tucker, M.A., Ord, T.J. and Rogers, T.L. (2014) ‘Evolutionary predictors of mammalian home range size: body mass, diet and the environment’, Global Ecology and Biogeography, 23(10), pp. 1105–1114. Available at: 10.1111/geb.12194.

Uezu, A., Metzger, J.P. and Vielliard, J.M.E. (2005) ‘Effects of structural and functional connectivity and patch size on the abundance of seven Atlantic Forest bird species’, Biological Conservation, 123(4), pp. 507–519. Available at: 10.1016/j.biocon.2005.01.001.

Valdés, A. et al. (2020) ‘High ecosystem service delivery potential of small woodlands in agricultural landscapes’, Journal of Applied Ecology, 57(1), pp. 4–16. Available at: 10.1111/1365-2664.13537.

Vannier, C., Bierry, A., et al. (2019) ‘Co-constructing future land-use scenarios for the Grenoble region, France’, Landscape and Urban Planning, 190, p. 103614. Available at: 10.1016/j.landurbplan.2019.103614.

Vannier, C., Lasseur, R., et al. (2019) ‘Mapping ecosystem services bundles in a heterogeneous mountain region’, Ecosystems and People, 15(1), pp. 74–88. Available at: 10.1080/26395916.2019.1570971.

Veldman, J.W. et al. (2019) ‘Comment on “The global tree restoration potential”’, Science, 366(6463), p. eaay7976. Available at: 10.1126/science.aay7976.

Vigiak, O. et al. (2003) ‘Spatial modeling of wind speed around windbreaks’, CATENA, 52(3), pp. 273–288. Available at: 10.1016/S0341-8162(03)00018-3.

Wartmann, F.M. et al. (2021) ‘Relating landscape ecological metrics with public survey data on perceived landscape quality and place attachment’, Landscape Ecology, 36(8), pp. 2367– 2393. Available at: 10.1007/s10980-021-01290-y.

Watts, K. et al. (2010) ‘Targeting and evaluating biodiversity conservation action within fragmented landscapes: an approach based on generic focal species and least-cost networks’, Landscape Ecology, 25(9), pp. 1305–1318. Available at: 10.1007/s10980-010-9507-9.

van der Zanden, E.H. et al. (2016) ‘Representing composition, spatial structure and management intensity of European agricultural landscapes: A new typology’, Landscape and Urban Planning, 150, pp. 36–49. Available at: 10.1016/j.landurbplan.2016.02.005.

Zoderer, B.M. et al. (2019) ‘Stakeholder perspectives on ecosystem service supply and ecosystem service demand bundles’, Ecosystem Services, 37, p. 100938. Available at: 10.1016/j.ecoser.2019.100938.

